# Psilocybin reduces heroin seeking behavior and modulates inflammatory gene expression in the nucleus accumbens and prefrontal cortex of male rats

**DOI:** 10.1101/2024.05.28.596205

**Authors:** Gabriele Floris, Konrad R. Dabrowski, Mary Tresa Zanda, Stephanie E. Daws

**Affiliations:** Center for Substance Abuse Research, Temple University, Philadelphia, PA USA; Department of Neural Sciences, Temple University, Philadelphia, PA USA; Department of Biology, Temple University, Philadelphia, PA USA

**Author notes:** Corresponding Author Contact Information: Stephanie Daws 3500 N Broad St MERB/ Rm 847 Philadelphia, PA 19140.

**Keywords:** opioid, serotonin, self-administration, relapse, drug-seeking, psychedelic, inflammation, cortex, nucleus accumbens, heroin, psilocybin, transcriptome

## Abstract

Preclinical and human studies indicate psilocybin may reduce perseverant maladaptive behaviors, including nicotine and alcohol seeking. Such studies in the opioid field are lacking, though opioids are involved in more >50% of overdose deaths. Psilocybin is an agonist at the serotonin 2A receptor (5-HT_2A_R), a well-documented target for modulation of drug seeking, and evidence suggests 5-HT_2A_R agonists may dampen motivation for opioids. We sought to investigate the therapeutic efficacy of psilocybin in mediating cessation of opioid use and maintenance of long-lasting abstinence from opioid seeking behavior in a rat model of heroin self-administration (SA). Psilocybin or 5-HT_2A_R antagonists ketanserin and volinanserin were administered systemically to rats prior to SA of 0.075 mg/kg/infusion of heroin, or relapse following forced abstinence. Psilocybin did not alter heroin taking, but a single exposure to 3.0 mg/kg psilocybin 4-24 hours prior to a relapse test blunted cue-induced heroin seeking. Conversely, 5-HT_2A_R antagonists exacerbated heroin relapse. To begin to elucidate mechanisms of psilocybin, drug-naïve rats received psilocybin and/or ketanserin, and tissue was collected from the prefrontal cortex (PFC), a region critical for drug seeking and responsive to psilocybin, 24 hours later for RNA-sequencing. 3.0 mg/kg psilocybin regulated ∼2-fold more genes in the PFC than 1.0 mg/kg, including genes involved in the cytoskeleton and cytokine signaling. Ketanserin blocked >90% of psilocybin-regulated genes, including the IL-17a cytokine receptor, *Il17ra*. Psychedelic compounds have reported anti-inflammatory properties, and therefore we performed a gene expression array to measure chemokine/cytokine molecules in the PFC of animals that displayed psilocybin-mediated inhibition of heroin seeking. Psilocybin regulated 4 genes, including *Il17a*, and a subset of genes correlated with relapse behavior. Selective inhibition of PFC IL-17a was sufficient to reduce heroin relapse. We conclude that psilocybin reduces heroin relapse and highlight IL-17a signaling as a potential downstream pathway of psilocybin that also reduces heroin seeking.

## Introduction

Opioid misuse contributed to more than half of the 100,000+ overdose deaths that occurred in the United States in 2021 ^1^. Elucidation of novel mechanisms that may reduce opioid use and sustain abstinence remains an essential and critical goal for the reduction of opioid-related misuse events. Clinical trials investigating the therapeutic efficacy of the psychedelic compound psilocybin for the treatment of neuropsychiatric disorders have shown promise for treating conditions plagued by perseverative maladaptive behaviors, including chronic drug seeking ^2–5^. Psilocybin and its active metabolite psilocin are serotonin 2A receptor (*Htr2a* [gene], 5-HT_2A_R [protein]) ^6–9^ agonists. It is well-established that pharmacological manipulation of 5-HT_2A_R can modulate drug self-administration (SA), reinstatement, and incubation of craving in preclinical models of psychostimulant, nicotine or alcohol seeking ^10–20^. Because psilocybin has demonstrated utility at reducing nicotine and alcohol use in humans ^2, 3^, it may also reduce seeking for other drugs, including opioids. In human subjects, the *HTR2A* gene was associated with heroin dependence, suggesting a link between 5-HT_2A_R and regulation of opioid seeking ^21, 22^. Several studies have investigated the impact of 5-HT_2A_R agonists on opioid seeking ^23–26^, but no studies to date have examined the impact of psilocybin on opioid seeking following forced abstinence. However, the limited data available demonstrates that 5-HT_2A_R agonists do not enhance opioid seeking and may decrease motivation for opioids ^23–26^. Thus, pharmacological tools that regulate 5-HT_2A_R signaling, such as psilocybin, may represent suitable candidates for the modulation of opioid seeking behavior.

Downstream of 5-HT_2A_R activation, molecular and cellular mechanisms of psilocybin may involve regulation of inflammatory signaling pathways because psychedelic compounds dampen activation of TNF-α processes ^27–29^. Transcriptome sequencing of postmortem brains following heroin overdose indicated that dysregulation of TNF-α/NF-kB-mediated inflammatory pathways in the nucleus accumbens (NAc) and prefrontal cortex (PFC), brain regions critical for drug reinforcement, is a major neuroadaptation following chronic heroin use ^30–32^. In this study, we aimed to address critical gaps in the field by investigating the putative efficacy of psilocybin in reducing opioid use and maintenance of abstinence from opioid seeking. We hypothesized that psilocybin would reduce heroin seeking and that antagonism of 5-HT_2A_R may have the opposite effect. We demonstrated that psilocybin reduces heroin seeking following forced abstinence and report the transcriptional effects of psilocybin on the PFC can be efficiently blocked with ketanserin. To delineate mechanisms of psilocybin on opioid seeking, we show that distinct patterns of inflammatory cytokine and chemokine genes are regulated in the PFC and NAc during psilocybin-mediated inhibition of heroin relapse and are associated with relapse behavior. Furthermore, we highlight the IL-17a pathway as a potential downstream pathway of psilocybin that is sufficient to reduce heroin relapse. Thus, we conclude that psilocybin reduces heroin relapse and regulates brain inflammatory gene expression, including components of the IL-17a pathway. Similarly, inhibition of PFC IL-17a with monoclonal antibodies also reduces heroin relapse. This suggests that diminishing this inflammatory pathway in the PFC may be one of the potential mechanisms through which psilocybin exerts its positive effects in reducing heroin-seeking behaviors.

## Methods and Materials

### Subjects

Adult male Sprague Dawley rats (Charles River Laboratories), aged 8 weeks on arrival, were used in this study. Rats for SA were kept on a reverse light/dark cycle (lights on at 9:00 p.m.; off at 9:00 a.m.) with constant room temperature (22 ± 2 °C) and humidity (40%). Rats were pair-housed upon arrival. Following intravenous catheter surgery, rats were single-housed for the remainder of the study. Food was provided *ad libitum,* except where described. All procedures were compliant with the National Institutes of Health’s Guide for the Care and Use of Laboratory Animals and approved by Temple University’s Institutional Animal Care and Use Committee.

### Drugs

Diamorphine hydrochloride (heroin) was provided by the National Institute on Drug Abuse drug supply program and dissolved in sterile saline (0.9% sodium chloride) for SA. Ketanserin (tartrate) (Cayman Chemical Company, Ann Arbor, MI; catalog # 22058) and volinanserin (Cayman Chemical Company, catalog #15936, also known as MDL 100907) were dissolved in 5% Tween- 80, then diluted with sterile saline. Psilocybin (Cayman Chemical Company; catalog # 14041) was dissolved in sterile saline. IL-17a monoclonal antibody (BioCell, Lebanon NH; catalog # BE0173) and IgG1 control (BioCell; catalog # BE0083) were diluted in *InVivo*Pure Dilution Buffer (BioCell; catalog # IP0080).

### Heroin self-administration (SA) and relapse

Jugular vein catheter implantation and long-access heroin SA were performed as previously described ^33, 34^, in 6-hour (hr) daily sessions on a fixed ratio (FR) 1 schedule at a dose of 0.075 mg/kg/infusion of heroin. Following SA, rats underwent forced abstinence in their home cage for 17 or 21 days (D). After abstinence, rats were re-exposed to SA chambers for one or three 30- minute relapse tests, during which all cues associated with active lever pressing were present, but no drug delivery occurred. Only one relapse test occurred per day, in consecutive days. Rats were immediately euthanized at the conclusion of behavioral assessments by rapid decapitation. Brains were removed, frozen by submersion in isopentane on a dry-ice platform and stored at - 80°C until dissection.

### Pharmacological manipulations

Psilocybin, ketanserin and volinanserin were administered prior to heroin SA on days 9-11, and prior to relapse tests following forced abstinence. For manipulations during SA and relapse, rats were balanced for drug intake after day 8 of SA and divided into experimental groups that were maintained throughout the duration of the study. For examination of the effects of a single dose of psilocybin on relapse, rats were divided into experimental groups based on heroin intake after day 10 of SA. 0.1, 1, or 3.0 mg/kg psilocybin, 0.1, 1, or 3.0 mg/kg ketanserin, or 0.04 or 0.2 mg/kg volinanserin were administered intraperitoneally (IP) either 15 minutes (ketanserin, volinanserin) or 4 hr (psilocybin) prior to the beginning of each SA or relapse session. One rat in the 1.0 mg/kg psilocybin group was excluded from the relapse test due to death during the abstinence period. For transcriptome profiling, drug-naïve rats were injected IP once with 1.0 mg/kg psilocybin, 3.0 mg/kg psilocybin, 3.0 mg/kg ketanserin, or 3.0 mg/kg of psilocybin and ketanserin simultaneously and euthanized 24 hr later. Sample sizes were as follows: psilocybin administration during SA and relapse- vehicle *n*=12; 0.1 mg/kg psilocybin *n*=5; 1.0 mg/kg psilocybin *n*=9; 3 mg/kg psilocybin *n*= 5; psilocybin administration once, prior to relapse- vehicle *n*=15; 3 mg/kg psilocybin, 4 hr *n*=8; 3 mg/kg psilocybin, 24 hr *n*=8; ketanserin administration during SA and relapse- vehicle *n*=12; 0.1 mg/kg ketanserin *n*=5; 1 mg/kg ketanserin *n*=8; 3 mg/kg ketanserin *n*=8; volinanserin administration during SA and relapse- vehicle *n*=6; 0.04 mg/kg volinanserin *n*=9; 0.2 mg/kg volinanserin *n*=10; For PFC infusion of IL-17a antibody prior to relapse- IgG *n*=8; IL-17a antibody *n*=7.

### Intra-PFC infusion of IL-17a antibody

Rats underwent stereotaxic surgery for the bilateral implantation of cannulas (Protech International, Inc, Boerne, TX), as we have described^35^, into the medial PFC, 5 days following the end of heroin SA, into the following coordinates from Bregma: anterior-posterior (AP) +2.7 mm, medial-lateral (ML) ±0.5 mm, and dorsal-ventral (DV) -3.5 mm. Rats received bilateral infusions of either 50 ng/ul IL-17a monoclonal antibody, or an IgG control, in a total volume of 1ul per hemisphere, delivered at a rate of 0.2 µL/min. The delivery of the correct volume was controlled using an infusion pump connected to two 10-microliter Hamilton syringes, which were linked to an infuser (Protech International, Inc,) via a tubing system. The first infusion was administered approximately 48 hours before the relapse test, and the second infusion occurred on the day of the relapse test, four hours prior. At the conclusion of the relapse test, rats were euthanized within one hour, their brains were rapidly removed and snap frozen by immersion in isopentane on a dry ice platform. Verification of cannula placements was performed through visual analysis during the tissue punching for the extraction of the PFC, utilizing the same coordinates as those used for cannulation.

### RNA extraction and gene expression analyses

PFC or NAc (core and shell) tissue from rat brains was dissected on a cold plate at -20°C. Total RNA was extracted using the miRNeasy Mini Kit (Qiagen, Hilden, Germany), following manufacturer’s instructions. The RT² Profiler™ Rat Inflammatory Cytokines & Receptors PCR Array (Qiagen) was used to measure inflammatory cytokines and chemokines in PFC and NAc tissue of rats that underwent heroin SA, forced abstinence and a single injection of 3.0 mg/kg psilocybin 4 or 24 hr prior to one 30-minute relapse test. For qPCR measurement of *Fos* in the PFC of rats that received IL-17a antibody prior to a heroin relapse test, RNA was reverse transcribed into cDNA using the Maxima Reverse Transcriptase kit (Thermo Fisher Scientific) and qPCR was performed using the PrimeTime Gene Expression Master Mix and gene assays (Integrated DNA Technologies, Coralville, IA). Primer sequences are available in the supplement. The RNA sequencing of PFC tissue from drug-naïve animals 24 hr following administration of psilocybin and/or ketanserin was performed by Novogene. Raw sequencing data are available in the Gene Expression Omnibus (Accession # GSE235175). For RNA sequencing, *n*=4 for all groups.

### Statistics

Statistical analyses were performed using GraphPad software (Prism version 10; GraphPad, San Diego, CA) and R. Error bars represent the mean ± the standard error of the mean (SEM). No randomization was performed in this study, due to the necessity to balance heroin intake between treatment groups. Sample sizes for experiments were determined based on similar studies in the neuroscience field. For behavioral data, repeated measures analysis of variance (RM-ANOVA) was used to analyze differences in lever pressing or infusions during SA. Outlier analysis was performed using the ROUT method in GraphPad. Shapiro-Wilk normality tests were performed to evaluate the distribution of data and similar levels of variation were observed between groups. For analysis of behavioral data at a single SA day or relapse test with a normal distribution, one-way ANOVAs were performed with post hoc Tukey tests to determine differences between more than two groups. For analysis of behavioral data at a single SA day or relapse test without a normal distribution, Kruskal-Wallis tests with post hoc Dunn’s tests were used to compare 3 or more groups. Paired t-tests were used to compare drug SA within individual rats before and after administration of psilocybin or ketanserin. Pearson correlations were used to compare the relationship between gene expression and relapse responses. For qPCR, differential expression analyses were performed using the ΔΔct method of analysis ^36^, followed by unpaired t-test between treatment groups. Replicates were excluded from qPCR analysis if a Ct value ≥ 35. For RNA-seq, DESeq2 R package (1.20.0) ^37^ was used to determine differential expression between two groups, and the GeneOverlap package in R Bioconductor, version 1.36.0 was used to determine statistical significance of two transcriptome datasets using the Fisher’s exact test. RNA sequencing was performed on biological replicate samples. Results from pharmacological manipulations were obtained from 2-3 cohorts of animals per manipulation. A p value of <0.05 was considered statistically significant.

**Additional methodological details are available in the supplemental material.**

## Results

### Psilocybin reduced heroin seeking, but not taking

To investigate the potential therapeutic efficacy of psilocybin in mediating cessation of opioid use and maintenance of long-lasting abstinence from opioid seeking behavior, we first tested the consequences of psilocybin on heroin SA (Figure 1A). Based on previous observations that 5-HT_2A_R agonists may decrease motivation for opioids^23, 24^, we hypothesized that psilocybin may reduce heroin taking. Adult male rats were trained to self-administer heroin in 6 hr daily sessions at a dose of 0.075mg/kg/infusion for 8 days (Figure 1B-F). Rats were separated into 4 treatment groups, with active lever responses and infusions during SA balanced: 0.1, 1, or 3.0 mg/kg psilocybin and saline vehicle (Figure 1D, F). On D9, 10 and 11 of heroin SA, rats received a single IP injection of psilocybin or vehicle 4 hr prior to SA. The 4 hr timepoint was chosen because at this timepoint, the psilocybin-induced head twitch response, thought to represent a hallucinogenic effect, has subsided and psilocybin will have been cleared from the brain and blood, thus eliminating any motor impairment confounds ^38–43^. Doses were chosen based on prior research involving psilocybin in rats ^18, 19, 41^. The 0.1 mg/kg dose is categorized as a microdose, given that it is one-tenth of the minimum dose required for eliciting behavioral and cellular effects in rats ^18, 44^. The 1 mg/kg dose is among the most extensively studied doses in rodents for both behavioral and molecular effects^44–46^, while 3 mg/kg falls within the range of therapeutic doses for humans ^18^. RM-ANOVA analysis revealed no main effect of psilocybin treatment or psilocybin by time interactions for any dose of psilocybin compared to vehicle for either lever presses or infusions (Figure 1G, H). Likewise, lever responses and infusions during D9-11 did not differ for SA of heroin when rats were pretreated with psilocybin. Analysis of individual rats’ heroin intake determined that rats do not significantly alter SA of heroin at the 0.075 mg/kg/infusion dose behavior following psilocybin (Supplemental Figure 1).

**Figure 1:**
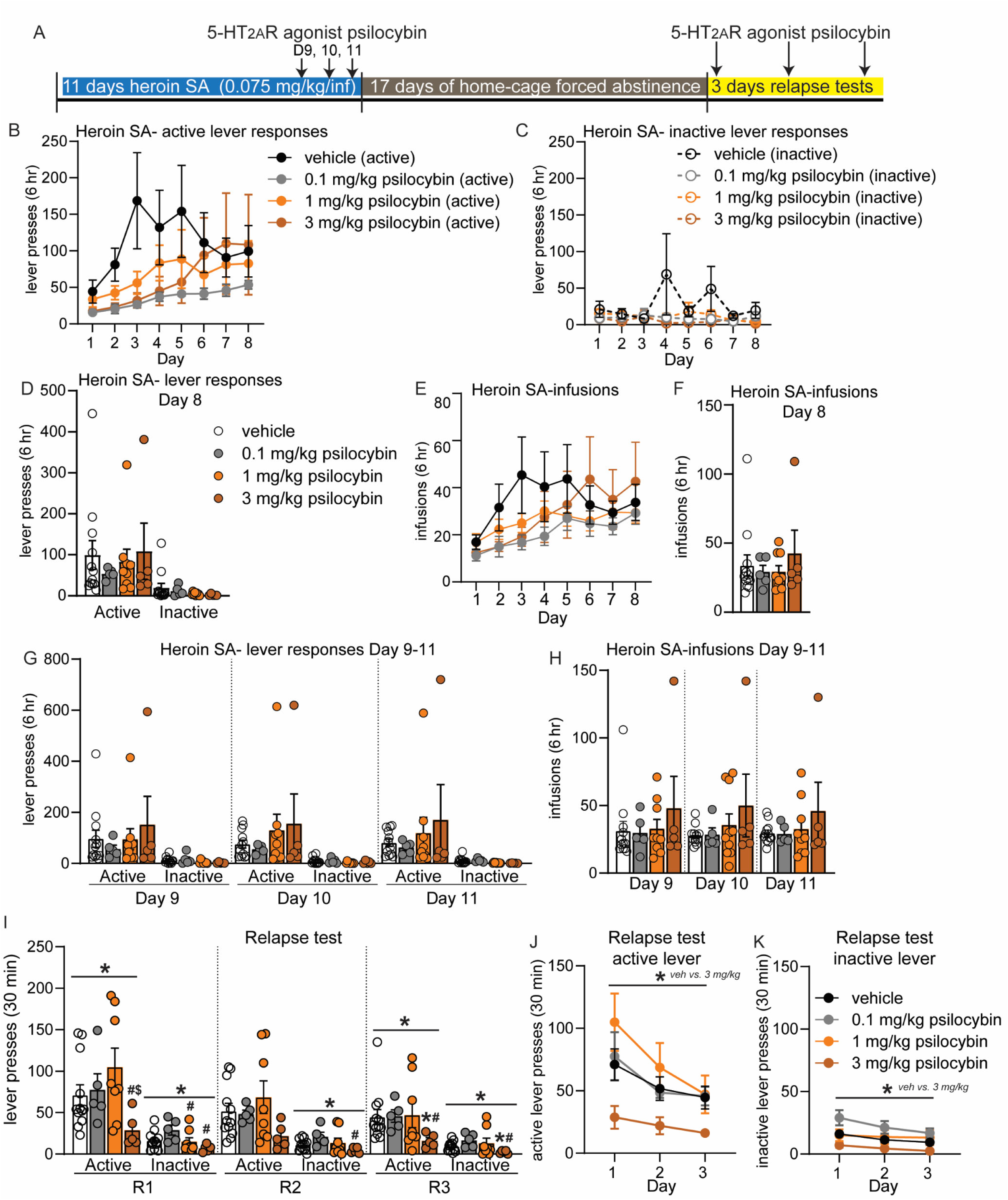
Psilocybin reduces heroin relapse after forced abstinence but not heroin intake. (A) Overview of experiment. Animals underwent heroin SA for 8 days. On days 9-11, animals were pretreated with 5-HT_2A_R agonist psilocybin 4 hr prior to SA. After 17D abstinence, animals underwent 3 relapse tests on consecutive days and were pretreated with psilocybin 4 hr prior to the test. (B-H) Shown are drug-paired active or inactive lever presses (B-D, G) and heroin infusions during SA (E, F, H). (B-F) SA behavior for each treatment group, prior to psilocybin treatment for days 1-8. Animals were split into treatment groups on day 8, with no significant differences observed in lever responses (D) or heroin infusions (F). (G-H) Heroin SA lever responses (G) and infusions (H) on days 9-11, following pretreatment with psilocybin. (I-K) Active and inactive lever presses during relapse tests on subsequent days (R1, R2, R3) for animals that were pretreated with psilocybin prior to the relapse test. (J-K) Shown are lever presses from panel I combined. Error bars indicate mean +/- standard error of the mean (S.E.M). *n*=5-12/group. Symbols above line indicate ANOVA: * p<0.05; Post hoc test indicated directly above individual histograms, * p<0.05 vs vehicle; # p<0.05 vs 0.1 mg/kg psilocybin; $ p<0.05 vs 1 mg/kg psilocybin.

We next examined the effect of psilocybin on heroin seeking and relapse after homecage forced abstinence (Figure 1A). Following 17D abstinence, rats were again injected with 0.1, 1 or 3 mg/kg psilocybin 4 hr prior to a 30-minute cue-induced relapse test in the SA chamber on three consecutive days (Figure 1A). Psilocybin pretreatment altered the response to heroin cues during each relapse test, with a significant reduction in active lever responses observed for rats treated with 3 mg/kg psilocybin on days 1 and 3 (Kruskal-Wallis test, R1: KW statistic = 8.33, p=0.04; R2: KW statistic = 5.65, p=0.130; R3: KW statistic = 7.85, p=0.049; Figure I). At the first relapse test, rats treated with 3.0 mg/kg psilocybin made significantly less active lever presses than 0.1 mg/kg and 1 mg/kg groups, with a trend for a reduction versus (vs) the vehicle group (Dunn’s tests, 3 mg/kg psilocybin vs vehicle: p=0.069; vs 0.1 mg/kg psilocybin: p=0.034; vs 1 mg/kg: p=0.005; Figure I). At the third relapse test, rats treated with 3 mg/kg psilocybin made significantly less active lever responses than vehicle and 0.1 mg/kg treated rats (Dunn’s tests, 3 mg/kg vs vehicle: p=0.016; vs 0.1 mg/kg psilocybin: p=0.010). Rats also differed in inactive lever responses when pretreated with psilocybin on all relapse test days (Kruskal-Wallis test, R1: KW statistic = 9.34, p=0.025; Dunn’s tests: 1 mg/kg vs 0.1 mg/kg psilocybin, p=0.034; 0.1 mg/kg vs 3 mg/kg psilocybin, p=0.003; R2: KW statistic = 7.96, p=0.047; Dunn’s tests: 0.1 mg/kg vs 3 mg/kg psilocybin, p=0.006; R3: KW statistic = 8.77, p=0.032; Dunn’s tests: 3 mg/kg psilocybin vs vehicle, p=0.031; 0.1 mg/kg vs 3 mg/kg psilocybin, p=0.004; Figure 2A). Rats treated with 0.1 mg/kg or 1.0 mg/kg psilocybin did not differ from vehicle. RM-ANOVA of lever responses for relapse days 1, 2, and 3 revealed a significant main effect of 3.0 mg/kg psilocybin treatment on both active and inactive lever presses, with psilocybin resulting in consistently lower lever responses (RM- ANOVA- vehicle vs 3 mg/kg psilocybin active lever: F(1, 15) = 5.60, p=0.032; inactive lever: F(1,15) = 9.373, p=0.008; Figure 1J, K). Thus, pretreatment with 3.0 mg/kg psilocybin diminishes heroin relapse after forced abstinence. This data suggests that activation of 5-HT_2A_R by 3.0 mg/kg psilocybin may be beneficial for reducing future heroin seeking behavior following abstinence.

**Figure 2:**
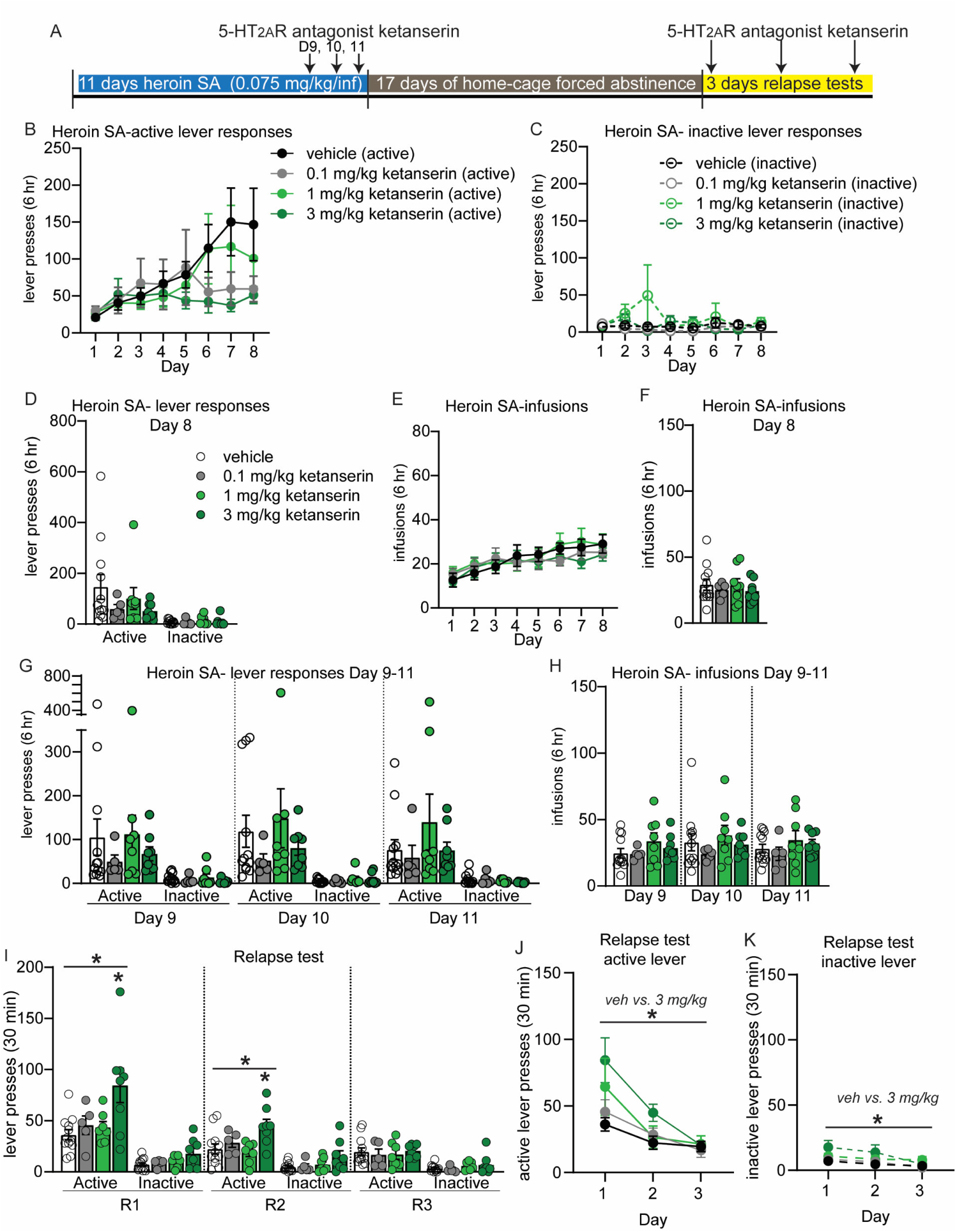
5-HT_2A_R antagonist ketanserin increases heroin self-administration and may exacerbate heroin relapse after forced abstinence at a dose of 3 mg/kg. (A) Overview of experiment. Animals underwent heroin SA for 8 days. On days 9-11, animals were pretreated with 5-HT_2A_R antagonist ketanserin IP 15 min prior to SA. After 17D abstinence, animals underwent 3 relapse tests on consecutive days and were pretreated with ketanserin 15 min prior to the test. (B-H) Shown are drug-paired active or inactive lever presses (B-D, G) and heroin infusions during SA (E, F, H). (B-F) SA behavior for each treatment group, prior to ketanserin treatment for days 1-8. Animals were split into treatment groups on day 8, with no significant differences observed in lever responses (D) or heroin infusions (F). (G-H) Heroin SA lever responses (G) and infusions (H) on days 9-11, following pretreatment with ketanserin. (I-K) Active and inactive lever presses during relapse tests on subsequent days (R1, R2, R3) for animals that were pretreated with ketanserin prior to the relapse test. (J, K) Shown are active lever presses from panel I combined. Error bars indicate mean +/- S.E.M. Post hoc test indicated directly above individual histograms, * p<0.05; *n*=5-12/group.

### 5-HT2AR antagonists may increase heroin intake and seeking

To provide more insight into the impact of 5-HT_2A_R signaling on heroin SA and relapse, we performed the same experiments with the widely-used 5-HT_2A_R antagonist ketanserin and the more selective antagonist volinanserin. Rats self-administered heroin for 8 days, then received 0.1, 1 or 3 mg/kg ketanserin 15 min prior to SA on D9-11 (Figure 2A). This timepoint was chosen to allow for comparisons to prior studies that have tested 5-HT_2A_R antagonists on drug seeking ^15, 17^. No differences in lever responses or heroin infusions were present prior to (Figure 2B-F) or following ketanserin, vs vehicle controls rats (Figure 3G, H). However, analysis of individual rats’ behavior during SA revealed a significant increase in heroin SA for rats in the 3.0 mg/kg ketanserin group (Supplemental Figure 2). Each rat pretreated with 3.0 mg/kg ketanserin prior to SA made more heroin infusions in the three days post treatment (9-11) compared to the three days preceding treatment (6-8) (Paired t-test 3.0 mg/kg ketanserin: t (7) =3.980, p=0.005; Supplemental Figure 2). Drug infusions for rats in the 3.0 mg/kg ketanserin group was steady throughout the 6 hr of SA, indicating ketanserin induced a consistent increase in heroin SA (Supplemental Figure 2). After 17D abstinence, rats received IP injection of 0.1, 1, or 3.0 mg/kg ketanserin 15 min prior to relapse tests on three consecutive days (Figure 2I-K). One-way ANOVA on active lever responses revealed a significant effect of ketanserin treatment on relapse days 1 and 2 (one-way ANOVA, R1: F (3, 28) = 5.116, p=.006; R2: F (3, 28) = 4.548, p=0.010; Figure 2I). Rats treated with 3.0 mg/kg ketanserin made more active lever presses compared to vehicle controls at the first two relapse tests (Tukey post hoc test, active lever vehicle vs 3.0 mg/kg ketanserin, R1: p=0.004; R2: p=0.017; Figure 2I). RM-ANOVA of lever pressing over the course of all three relapse tests revealed a significant main effect of 3 mg/kg ketanserin, as well as a significant time X 3 mg/kg ketanserin treatment interaction for active lever responses, and a significant main effect of 3 mg/kg ketanserin treatment for inactive lever responses (RM-ANOVA, active lever responses R1-3, 3 mg/kg ketanserin treatment effect: F (1, 18) = 8.333, p=0.010; time X treatment effect, active lever: F(2,36) = 10.67, p=0.0002; main effect of ketanserin treatment for inactive lever: F(1, 18) = 7.134, p=0.016; Figure 2J, K). Thus, ketanserin may exacerbate heroin seeking at the 3.0 mg/kg dose, which is in alignment with the increase in lever pressing and infusions observed during heroin SA. Importantly, we verified that administration of ketanserin or psilocybin at the time points tested did not induce overall gross changes in locomotor behavior (Supplemental Figure 3).

**Figure 3:**
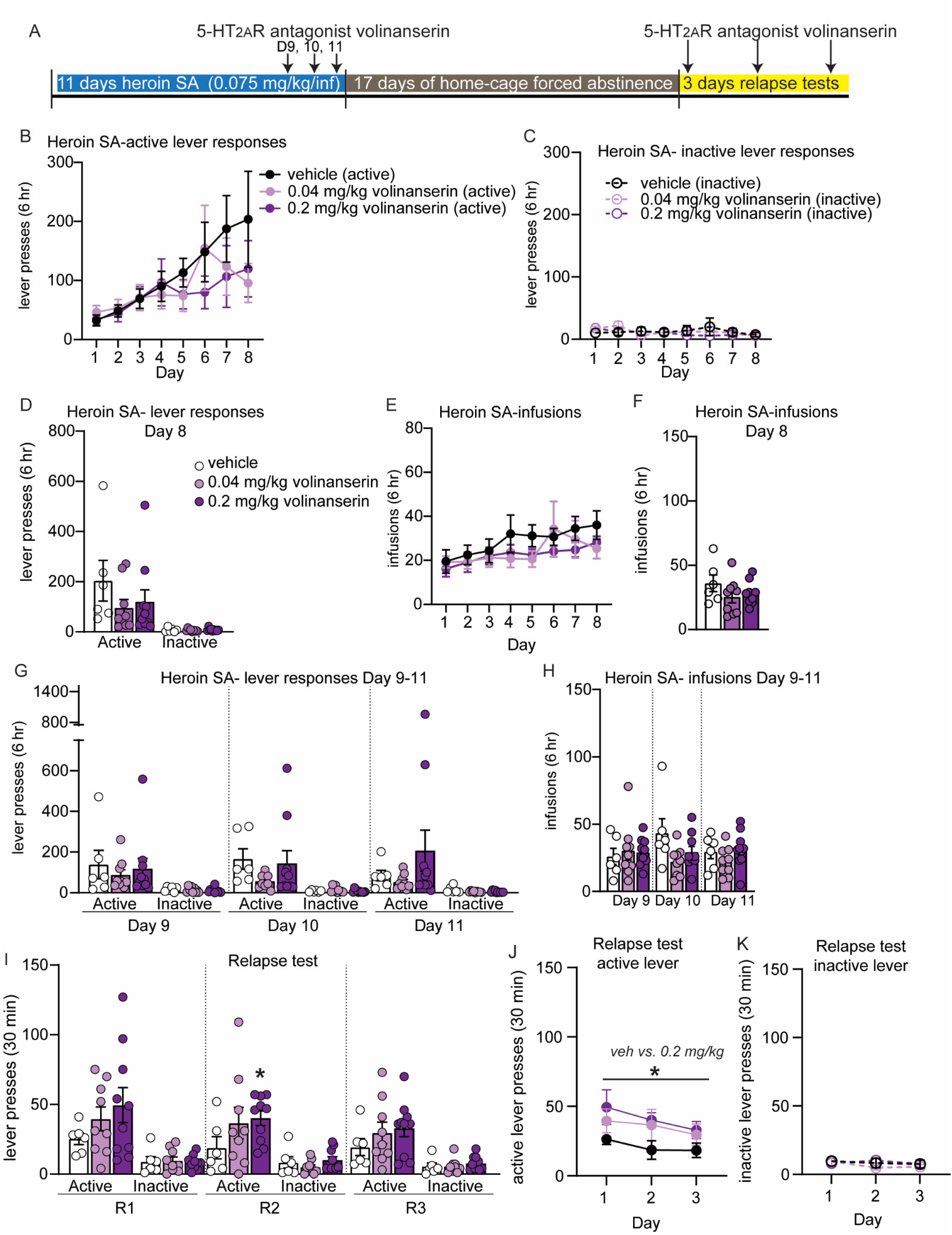
The selective 5-HT_2A_R antagonist volinanserin increases heroin relapse. (A) Overview of experiment. Animals underwent heroin SA for 8 days. On days 9-11, animals were pretreated with 5-HT_2A_R antagonist volinanserin IP 15 min prior to SA. After 17D abstinence, animals underwent 3 relapse tests on consecutive days and were pretreated with volinanserin 15 min prior to the test. (B-H) Shown are drug-paired active or inactive lever presses (B-D, G) and heroin infusions during SA (E, F, H). (B-F) SA behavior for each treatment group, prior to volinanserin treatment for days 1-8. Animals were split into treatment groups on day 8, with no significant differences observed in lever responses (D) or heroin infusions (F). (G- H) Heroin SA lever responses (G) and infusions (H) on days 9-11, following pretreatment with volinanserin. (I-K) Active and inactive lever presses during relapse tests on subsequent days (R1, R2, R3) for animals that were pretreated with volinanserin prior to the relapse test. (J, K) Shown are active lever presses from panel I combined. Error bars indicate mean +/- S.E.M. * above solid line denotes p<0.05 for RM-ANOVA between .2 mg/kg volinanserin and vehicle; * directly above histogram indicates unpaired t-test between 0.2 mg/kg volinanserin and vehicle with p<0.05. *n*=6-10/group.

Ketanserin has been widely used in behavioral pharmacology as a 5-HT_2A_R antagonist, despite the fact that it also has some activity at 5-HT_2C_R. Given this fact, coupled with the published studies demonstrating that 5-HT_2C_R antagonism may be beneficial for reducing drug seeking, we chose to repeat our studies using the more selective 5-HT_2A_R antagonist volinanserin. Rats underwent identical procedures, as described above for ketanserin, except were treated with either 0.04 or 0.2 mg/kg volinanserin or vehicles, 15 min prior to each behavior examined. Similar to our reported findings with psilocybin and ketanserin, volinanserin-treated rats did not differ in their heroin SA compared to vehicle-treated control rats (Figure 3B-H). Rats underwent 17D forced abstinence and were then treated with volinanserin or vehicle 15 min prior to the cue-induced relapse test. One-way ANOVA of active and inactive lever responses during each relapse day did not yield statistical significance. However, a trend for an increase in active lever responses was observed for rats that received 0.2 mg/kg volinanserin compared to vehicle-treated rats, with a significant increase in active lever responses observed on relapse day 2 (unpaired t-test, 0.2 mg/kg volinanserin vs vehicle, t(14) = 2.352, p=0.034; Fig. 3I). RM-ANOVA of lever responses over all three days of relapse tests revealed a significant main effect of 0.2 mg/kg volinanserin treatment compared to vehicle treated rats, with this dose of volinanserin significantly elevating active, but not inactive, lever responses (RM-ANOVA, F(1, 15) = 5.575, p=0.0322; Figure 3J, K). These data lead us to conclude that antagonism of 5-HT_2A_R prior to cue-induced relapse increases heroin seeking behavior.

### A single exposure to psilocybin alters the PFC transcriptome

To discern a molecular mechanism for psilocybin-mediated inhibition of heroin seeking, we measured mRNA expression of genes in the PFC (prelimbic and infralimbic medial prefrontal cortex) following acute exposure to psilocybin. The PFC mediates drug seeking behavior ^32, 47, 48^, and psilocybin impacts PFC circuit activation ^45, 49, 50^. Given the observed reduction in heroin-seeking behavior after acute 3.0 mg/kg psilocybin treatment but not 1 mg/kg, we decided to examine psilocybin-induced transcriptional changes that occur in the PFC following the 3 mg/kg dose. RNA sequencing was performed on PFC tissue from drug-naïve rats that received IP injection of 1.0 mg/kg psilocybin + vehicle, 3.0 mg/kg psilocybin + vehicle, 3.0 mg/kg ketanserin + vehicle, 3.0 mg/kg psilocybin + 3.0 mg/kg ketanserin, or both vehicles alone (Figure 4 and Supplemental Excel Tables SE1-8). Each animal received two injections to account for exposure to psilocybin (saline) or ketanserin (5% tween-80 in saline) vehicles. The choice to use drug-naïve rats was made to eliminate confounding factors related to chronic heroin exposure and withdrawal periods, ensuring a clear setting for pharmacological comparison. This setting allowed for an unambiguous evaluation of the effects of the 3 mg psilocybin treatment against the non-effective 1 mg dose and the 3 mg ketanserin treatment, which produced opposite results. 3.0 mg/kg psilocybin regulated expression of 314 genes in the PFC, while ketanserin regulated 214 (Figure 4B-C and Supplemental excel tables SE1-4). Coadministration of 3.0 mg/kg ketanserin/psilocybin blocked regulation of >90% of psilocybin-regulated genes in the PFC, while 1.0 mg/kg psilocybin regulated expression of only 177 genes (Figure 4D-E). Venn diagrams of genes regulated in more than one condition indicated that coadministration of 3.0 mg/kg ketanserin/psilocybin blocked ketanserin-specific or psilocybin-specific genes, but 90 genes were uniquely regulated by the combination of the two drugs (Figure 4F). An alluvial plot demonstrated that most genes regulated by 1.0 or 3.0 mg/kg psilocybin were muted by dual application of 3.0 mg/kg psilocybin/ketanserin or 3.0 mg/kg ketanserin alone (Supplemental Figure 4 and Supplemental Excel Tables SE6-8). While a quarter of genes were regulated by both 1.0 and 3.0 mg/kg psilocybin, the two doses resulted in unique in molecular consequences. We performed rank-rank hypergeometric overlap (RRHO) analysis ^51^ and observed coordinated gene expression between the two doses of psilocybin (Figure 4G and Supplemental Excel Table SE5). Conversely, an RRHO analysis of 3.0 mg/kg psilocybin vs 3.0 mg/kg ketanserin or 3.0 mg/kg psilocybin/ketanserin vs 3.0 mg/kg psilocybin alone revealed that treatment with ketanserin blocks PFC gene expression induced by psilocybin (Figure 4H-I). Interestingly RRHO analysis of 3.0 mg/kg psilocybin/ketanserin vs 3.0 mg/kg ketanserin alone suggested that psilocybin may not combat many of the effects of ketanserin (Figure 4J).

**Figure 4:**
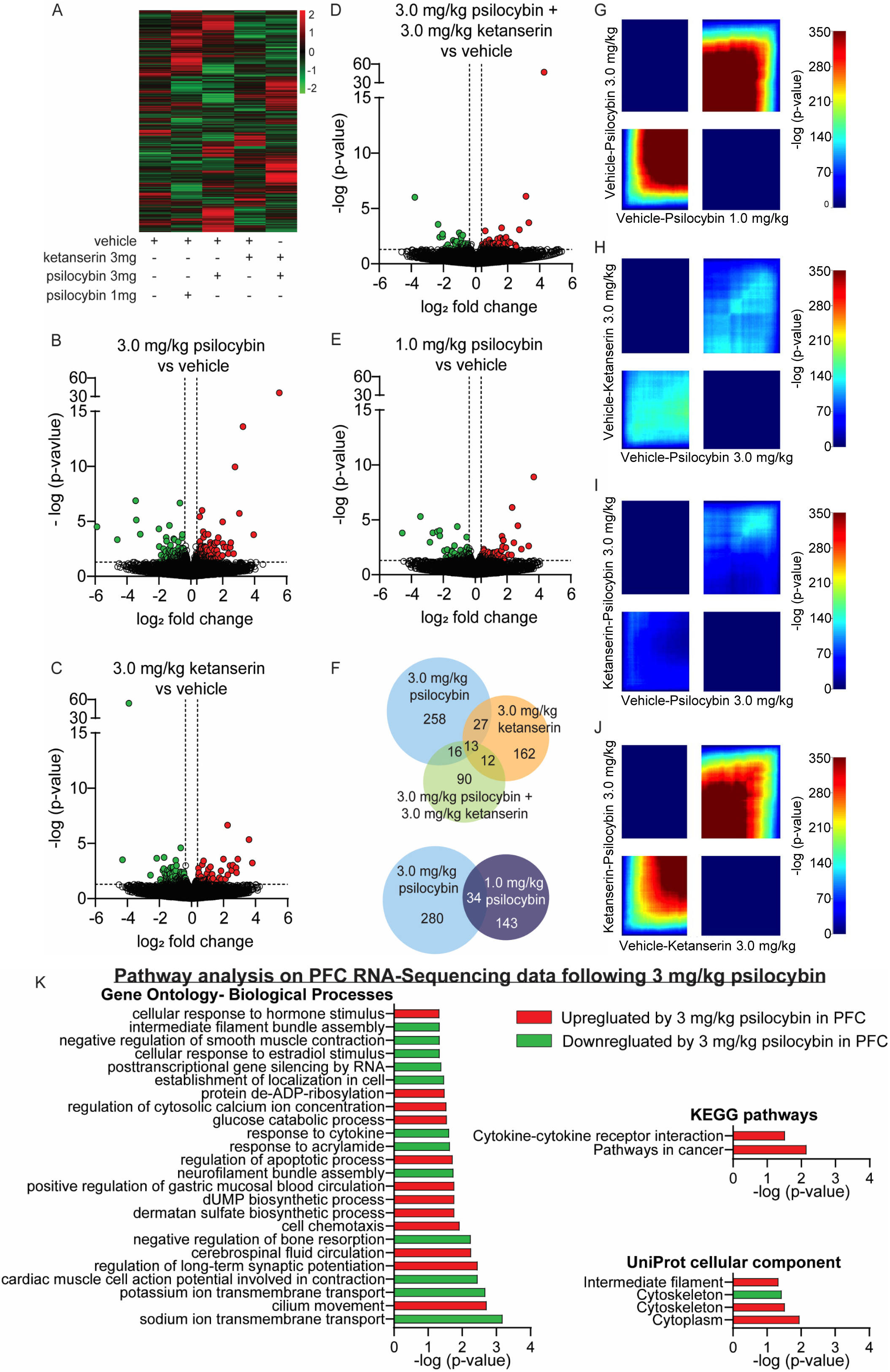
Psilocybin induces unique gene expression profiles in the PFC at varying doses that are blocked by ketanserin. (A) Heatmap of PFC gene expression profiles obtained from tissue collected 24 hours after a single injection of psilocybin and/or ketanserin. (B-E) Volcano plots depicting differential gene expression in the PFC between vehicle treated animals and treatment with 3.0 mg/kg psilocybin (B), 3.0 mg/kg ketanserin (C), 3.0 mg/kg psilocybin and ketanserin (D), or 1.0 mg/kg psilocybin (E). (F) Venn diagram of total genes regulated by psilocybin and/or ketanserin in the PFC and the number of genes regulated by more than one treatment group. (G-H) RRHO plots were generated to compare all gene expression data between animals treated with 3.0 mg/kg psilocybin and 1.0 mg/kg psilocybin (G), 3.0 mg/kg ketanserin and 3.0 mg/kg psilocybin (H), 3.0 mg/kg ketanserin + 3.0 mg/kg psilocybin and psilocybin 3mg/kg alone (I), or 3.0 mg/kg ketanserin + 3.0 mg/kg psilocybin and ketanserin 3mg/kg alone (J). Lower left quadrant represents genes up-regulated in both groups while upper right quadrant represents genes down-regulated in both groups. (K) Gene ontology, KEGG pathway analysis and Uniprot cellular compartments of genes regulated in the PFC following a single exposure of 3.0 mg/kg psilocybin in drug-naïve animals. *n*=4/group.

Pathway analysis and gene ontology assessment of genes significantly altered in the PFC following acute psilocybin revealed psilocybin-regulated genes belonged to gene ontology terms that included ‘sodium ion transmembrane transport,’ ‘regulation of long-term synaptic potentiation,’ ‘neurofilament bundle assembly,’ and ‘response to cytokine’ (Figure 4K). However, only two KEGG pathways were significantly regulated by psilocybin, ‘Cytokine-cytokine receptor interaction’ and ‘Pathways in cancer.’ Assessment of the Uniprot cellular component of the psilocybin-regulated genes revealed significant enrichment for terms related to ‘Cytoplasm,’ ‘Cytoskeleton’ and ‘Intermediate filament.’

### Pre-treatment with a single dose of psilocybin reduced cue-induced heroin seeking

Psilocybin-mediated reduction of heroin seeking at the 3.0 mg/kg dosage (Figure 1I-K) represents an intriguing finding, as psilocybin has been proposed as a potentially effective, low-risk, long-lasting therapeutic tool for treating patients with substance use disorders (SUD) ^52–55^ and just one exposure to psilocybin has long-lasting behavioral and molecular consequences ^56^. To further explore the impact of psilocybin on heroin seeking in another translationally-relevant administration protocol, we evaluated the temporal aspect of a single psilocybin exposure on heroin relapse by treating additional groups of rats with 3.0 mg/kg psilocybin once, either 4 or 24 hr prior to a relapse test (Figure 5A). All rats acquired heroin SA over 10 days (Figure 5B-C) and were divided into experimental groups after SA, with lever response and heroin intake during SA balanced (Figure 5D-E). After 21D abstinence, rats were injected IP with 3.0 mg/kg psilocybin or saline vehicle either 4 hr or 24 hr prior to a single 30-minute relapse test. Vehicle-treated animals at each timepoint performed almost identically in both SA as well as the relapse test, and were therefore collapsed into a single vehicle group (Supplemental Figure 5; relapse active lever presses for vehicle 4 hr = 77.14 +/- 11.21; vehicle 24 hr= 76.25 +/- 15.73). One-way ANOVA analysis of active lever response during the relapse test revealed that a single exposure to 3 mg/kg psilocybin, either 4 or 24 hours prior, was sufficient to significantly reduce cue-induced heroin seeking behavior following forced abstinence (Kruskal-Wallis test: KW statistic = 7.657, p=0.025; Dunn’s tests: vehicle vs 3 mg/kg psilocybin 4 hr, p=0.0379; vehicle vs 3 mg/kg psilocybin 24 hr, p=0.0148; Figure 5G). No significant differences between treatment groups were observed for inactive lever responses during the relapse test.

**Figure 5:**
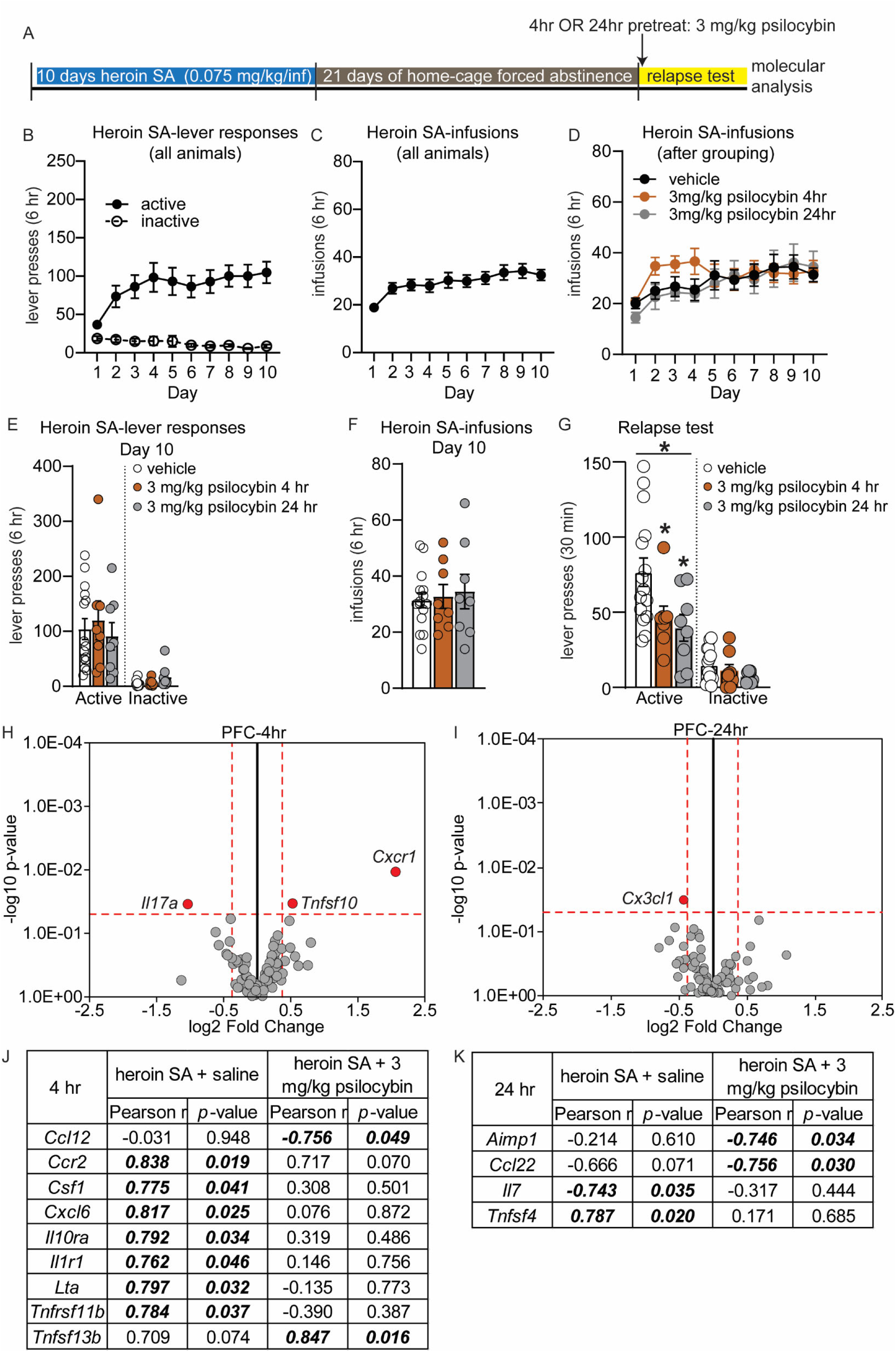
Psilocybin pretreatment blunts heroin relapse and regulates gene expression of inflammatory cytokine and chemokines in the PFC during inhibition of heroin seeking. (A) Overview of experiment. Animals underwent heroin SA for 10 days, 21D forced abstinence and then a 30-minute relapse test. Animals were pretreated with a single dose of 3.0 mg/kg psilocybin or saline vehicle either 4 or 24 hr prior to the test. Animals were euthanized within 1 hour of completion of the relapse test. (A-F) Shown are drug-paired active or inactive lever presses (B, E) and heroin infusions during SA (C, D, F) for each group, prior to any psilocybin treatment. Animals were balanced for lever responses and heroin infusions (D-F) on day 10 of SA. (G) Active and inactive lever presses during the relapse test for rats that received 3 mg/kg psilocybin 4 or 24 hr prior to testing. <u>Asterisk above solid line</u> denotes p<0.05 for Kruskal-Wallis test; * directly above histogram denotes p<0.05 vs vehicle for Post hoc test. Error bars indicate mean +/- S.E.M. *n*=8-15/group. (H-I) Volcano plot depicting inflammatory cytokine and chemokine genes, obtained by qPCR array, that were differentially expressed in the PFC of rats that were treated with 3.0 mg/kg psilocybin 4 hr (H) or 24 hr (I) prior to a relapse test, vs vehicle treated animals. Horizontal dotted line denotes p value of 0.05. Vertical dotted line denotes fold change of +/- 30%. Grey circles represent genes that were not significantly altered. Red dots represent genes that meet criteria for significance (J-K) Results of correlation analysis of relapse behavior of rats (from G) with inflammatory cytokine or chemokine gene expression data for the PFC that had psilocybin 4 hr (J) or 24 hr (K) prior to a relapse test. Italicized, bolded values were statistically significant.

### Psilocybin-mediated inhibition of heroin seeking is accompanied by modulation of inflammatory chemokines and cytokines in the PFC and NAc

The gene ontology and KEGG pathway analysis overlapped to indicate that psilocybin alters PFC expression of genes related to cytokines signaling pathways. We focused further on psilocybin-induced regulation of inflammatory molecules because activation of 5-HT_2A_R by agonists produces anti-inflammatory effects; cytokine and chemokine signaling is involved in the rewarding effects of drugs; and human transcriptomics data demonstrated heroin use results in dysregulation of PFC and NAc inflammatory pathways ^29, 30, 57, 58^. A focused qPCR array on 84 inflammatory cytokines and chemokines was performed using PFC tissue from rats that underwent the relapse test in Figure 6G after either 4 or 24 hr pretreatment with 3.0 mg/kg psilocybin (Figure 5H-I). In the PFC of rats pretreated 4 hr before the relapse test with psilocybin, psilocybin-mediated inhibition of heroin relapse was accompanied by significant downregulation of Interleukin 17A (*Il17a*), and upregulation of C-X-C Motif Chemokine Receptor 1 (*Cxcr1*) and TNF Superfamily Member 10 (*Tnfsf10*) (unpaired t-test, vehicle vs 3.0 mg/kg psilocybin: *Il17a*, t(12) = 2.39, p=0.034; *Cxcr1*, t(12) = 3.02, p=0.011; *Tnfsf10*, t(12) = 2.40, p=0.036; Figure 5H). Interestingly, the *Il17a* receptor, *Il17ra*, was among the most significantly downregulated genes in the PFC in our transcriptome analysis following 3 mg/kg psilocybin (Supplemental excel file). In animals pretreated 24 hr before the relapse test with psilocybin, psilocybin-mediated inhibition of heroin seeking was associated with significant downregulation of C-X3-C Motif Chemokine Ligand 1 (*Cx3cl1*) in the PFC (unpaired t-test, vehicle vs. 3.0 mg/kg psilocybin: t(14)= 2.37, p=0.032; Figure 5I).

**Figure 6:**
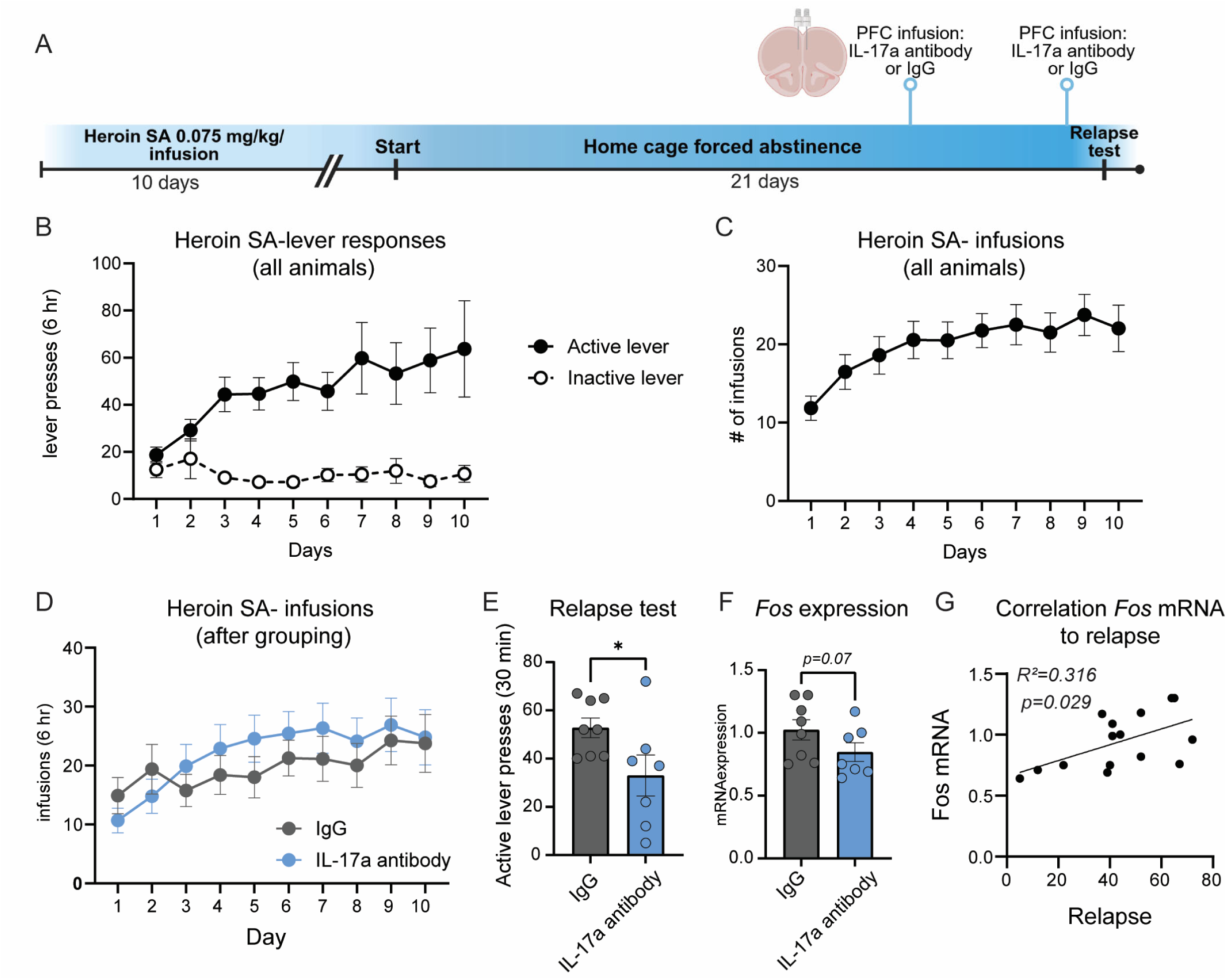
Selective inhibition of IL-17a signaling in the PFC is sufficient to reduce heroin seeking following forced abstinence. (A) Experimental overview: rats underwent jugular vein catheterization and implantation of PFC cannula. Following recovery, rats underwent heroin SA, forced abstinence and a relapse test. 72 and 24 hours prior to the relapse test, rats received infusion of 50 ng/hemisphere of IL-17a antibody or an IgG control directly into the PFC. (B) Active and inactive lever responses during heroin SA, prior to PFC infusion of IL-17a antibody. (C, D) Heroin infusions made during heroin SA for all rats (C) and depiction of infusions for each treatment group after regrouping into balanced treatment groups (D). (E) Active lever presses during a 30 min relapse test for rats that received IL-17a antibody into the PFC and IgG controls. (F) *Fos* mRNA in the PFC of rats that received IL-17a antibody prior to a relapse test. (G) Pearson correlation of *Fos* mRNA expression with active lever responses during the relapse test for rats that received IL-17a antibody or IgG into the PFC prior to the relapse test. Error +/- S.E.M. *p<0.05. *n*=7-8/group.

Correlation analysis of relapse performance with gene expression revealed 7 genes were positively correlated with active lever responses in the PFC of heroin SA animals pretreated with saline, but not psilocybin 4 hr prior: C-C Motif Chemokine Receptor 2 (*Ccr2*), Colony Stimulating Factor 1 (*Csf1*), C-X-C Motif Chemokine Ligand 6 (*Cxcl6*), Interleukin 10 Receptor Subunit Alpha (*Il10ra*), Interleukin 1 Receptor Type 1 (*Il1r1*), Lymphotoxin Alpha (*Lta*) and *Tnfrsf11b* (Figure 5J). Conversely, expression of C-C motif chemokine ligand 12 (*Ccl12*) was negatively correlated with relapse in rats that were pretreated 4 hr prior with psilocybin, but not saline, while expression of TNF Superfamily Member 13b (*Tnfsf13b*) was positively correlated with relapse only in psilocybin-treated rats (Figure 5J). Following the 24 hr pretreatment with psilocybin, significant negative correlation was observed between relapse behavior and *Ccl22*, as well as Aminoacyl TRNA Synthetase Complex Interacting Multifunctional Protein 1 (*Aimp1*) (Figure 5K).

We conducted further analysis in the NAc, another region sensitive to the modulatory action of inflammatory factors in drug-seeking behaviors^59, 60^, of animals that had completed the relapse test 4 hours following the acute administration of 3.0 mg/kg psilocybin to determine the specificity of cytokine and chemokine regulation following psilocybin-mediated inhibition of heroin seeking. In the NAc, psilocybin pre-treatment prior to relapse tests upregulated 4 genes: Chemokine (C-C motif) receptor 3 (*Ccr3*), Chemokine (C-X-C motif) receptor 3 (*Cxcr3*), Interleukin 1 beta (*Il1b*), and Tumor necrosis factor receptor superfamily, member 11b (*Tnfrsf11b*) (unpaired t-test, vehicle vs 3.0 mg/kg psilocybin: *Ccr3*, t(12) = 2.56, p=0.025; *Cxcr3*, t(12) = 2.45, p=0.031; *Il1b*, t(12) = 3.12, p=0.009; *Tnfrsf11b*, t(12) = 3.92, p=0.002; Supplemental Figure 6A). In the NAc, 7 genes were negatively correlated with relapse behavior in rats pretreated with psilocybin, but not saline: C-C Motif Chemokine Ligand 7 (*Ccl7*), C-C Motif Chemokine Receptor 10 (*Ccr10*), *Ccr2*, Fas Ligand (*Faslg*), Interleukin 33 (*Il33*), Platelet Factor 4 (*Pf4*), and *Tnfrsf11b* (Supplemental Figure 6B). These results demonstrate that psilocybin-mediated inhibition of heroin seeking is accompanied by regulation of inflammatory cytokines and chemokines in the NAc and PFC.

### Selective inhibition of Il17a in the PFC is sufficient to reduce heroin relapse

In multiple molecular analyses, we identified regulation of the IL-17 signaling pathway following psilocybin: 3 mg/kg psilocybin reduced *Il17ra*, the receptor for IL-17a, 24 hr later in the PFC; and pretreatment with 3 mg/kg psilocybin prior a relapse test reduced *Il17a* in the PFC during psilocybin-mediated inhibition of heroin relapse. Based on these data, we hypothesized that IL-17a signaling may function downstream of psilocybin to reduce heroin seeking behavior. To begin to explore this hypothesis, we selectively reduced IL-17a signaling in the PFC during heroin relapse. Rats were implanted with jugular vein catheters, then underwent heroin SA for 10D and forced abstinence for 21D (Figure 6A). Rats were grouped into IL-17a antibody or IgG control groups based on their SA behavior and did not differ in drug infusions or lever presses (Figure 6B-D). During abstinence, PFC cannula were implanted. 48 and 4 hr prior to a 30 min cue-induced relapse test, rats received infusion of 50 ng/hemisphere of IL-17a antibody or IgG control directly into the PFC (Fig. 6A). Selective infusion of IL-17a antibody into the PFC significantly reduced heroin seeking behavior during the relapse test (unpaired t-test, t(13) = 2.187, p=0.048; Fig. 6E). Rat brains were collected within 90 min of the relapse test and *Fos* mRNA expression was measured in the PFC, as a measure of activity. Rats infused with the IL- 17a antibody into the PFC displayed a trend for reduced *Fos* activity in the PFC that did not reach statistical significance (Mann-Whitney test U= 12.50, p=0.07; Fig. 6F). However, a correlation analysis revealed a significant negative correlation between *Fos* mRNA expression in the PFC following IL-17a antibody infusion and active lever responses during the relapse test (Pearson correlation: R^2^=0.316, p=0.0291; Fig. 6G). These data indicate that inhibiting PFC IL- 17a with monoclonal antibodies reduces heroin relapse. This reduction in relapse is correlated with decreased *Fos* mRNA in the PFC, suggesting lower PFC reactivity during heroin seeking after forced abstinence.

## Discussion

This study is the first to demonstrate that psilocybin reduces heroin seeking in a model of forced abstinence. Although we also pretreated rats with psilocybin during SA and did not observe a regulation of heroin intake, we excluded contribution of prior psilocybin exposure during SA to the relapse test results by replicating our findings of psilocybin-mediated inhibition of heroin seeking after a single injection of 3.0 mg/kg psilocybin only prior to the relapse test. Additionally, our study design allowed us to evaluate the effect of repeated psilocybin administration on heroin relapse, in comparison to acute psilocybin administration. Both administration paradigms are translationally relevant, as psilocybin can induce long-lasting changes in behavior and plasticity following a single exposure^56^, yet reports of ‘microdosing’, which has not been examined in models of drug seeking, may have therapeutic value for drug seeking behavior by protecting against stress-induced phenotypes^61^, such as stress-induced relapse.

We examined three doses of psilocybin and observed a pattern of decreased heroin seeking only in rats treated with 3.0 mg/kg. Lack of efficacy from the 1.0 mg/kg psilocybin dose on heroin relapse may be due to the fact that occupancy of 5-HT_2A_R is estimated to only be ∼30% with 1.0 mg/kg IP dosages ^62^. The 1 mg/kg dose is among the most extensively studied doses in rodents for both behavioral and molecular effects ^44–46^, while 3 mg/kg falls within the range of therapeutic doses for humans ^18^. While psilocybin does induce head-twitch responses in rodents, the 3 mg/kg dose has been reported to induce less head-twitch responses compared to the 1 mg/kg dose over time but 3 mg/kg induces a rapid, short-lived head-twitch response in rodents that may be further indicative of unique molecular and behavioral consequences that arise from the two doses ^39, 40^. Accordingly, our analyses revealed that the transcriptional consequences were distinct between the 1.0 and 3.0 mg/kg doses of psilocybin. In our study, rats underwent three relapse tests following 21D forced abstinence from heroin SA and we observed a rapid reduction of heroin seeking behavior in rats that received a pretreatment with 3 mg/kg psilocybin prior to the relapse test. The 30-minute relapse test functions similarly to extinction training, due to the fact that rats receive all drug-associated cues in the same context that they previously self-administered heroin. Therefore, the reduction of active lever responses in rats treated with 3 mg/kg psilocybin prior to the relapse test suggests that psilocybin may also facilitate extinction of opioid-associated drug memories. Conversely, ketanserin and volinanserin may delay extinction. The repeated relapse testing in Figures 1 and 2 highlight the potential for psilocybin to be used in combination with behavioral therapy to rapidly reduce drug seeking behavior and facilitate extinction of drug seeking.

Our observation of psilocybin-mediated inhibition of heroin seeking is in alignment with published studies that demonstrated that the 5-HT_2A_R psychedelic agonist 2,5-dimethoxy-4- iodoamphetamine (DOI) decreased demand for fentanyl and a non-hallucinogenic 5-HT_2A_R agonist tabernanthalog (TBG) reduced both heroin SA and seeking behavior ^23, 25, 26^. In contrast, a similar dose of 2.5 mg/kg psilocybin had no enduring impact on re-initiation of alcohol taking ^18^. This divergence likely arises from distinct methodological approaches, as we utilized a cue-induced relapse protocol without reintroduction of heroin, and administered a single psilocybin dose, as opposed to their two-dose regimen spaced a week apart. Subsequent investigation into the impact of psilocybin on alcohol SA demonstrated that a single psilocybin injection 4 hr before the relapse test effectively reduced alcohol SA ^19^. This suggests that the efficacy of psilocybin in reducing relapse may be contingent upon experimental conditions, dosing strategies, and drug re-exposure. Conversely, nicotine and cocaine studies demonstrate that inhibition of 5-HT_2A_R generally reduces seeking behavior, which suggests divergence of 5- HT_2A_R signaling with different drug classes ^10–12, 15–17^. However, human studies indicate that psilocybin may reduce nicotine craving ^3^, and therefore, it is possible that psilocybin may have therapeutic efficacy through non-5-HT_2A_R-mechanisms ^63^, or that humans have unique psychedelic experiences from psilocybin that may not be recapitulated in all rodent models. Recent studies suggest that the therapeutic effects of serotonergic psychedelics may be dissociable from their hallucinogenic effects^45, 64^. Specifically, it has been observed that while pretreatment with ketanserin can block the acute effects of psilocybin, the plasticity-promoting and antianhedonic effects of psilocybin were not reversed with ketanserin^64^. However, a recent study by Vargas et al.^65^ has revealed that the effects of psychedelics on synaptic plasticity can be inhibited by ketanserin at a dose ten times higher than the dose of psychedelic used. This finding suggests that additional research using 5-HT_2A_R knockout animals and/or different doses/timing of ketanserin administration in models of opioid SA may be necessary to further explore this significant, yet open question. However, it is important to note that the dose of psilocybin (3 mg/kg) that reduced heroin seeking in our model is higher than the 1 mg/kg dose of psilocybin used in the aforementioned studies that was sufficient for antidepressant and plasticity-promoting effects. Therefore, it is challenging to draw conclusions from these discrepancies, and it is possible that different mechanisms may underlie the antidepressant and anti-drug seeking effects. In our study, we offer a different perspective, demonstrating that the transcriptional effects of psilocybin are largely, though not completely, inhibited by ketanserin administration. Hence, the current study, together with cumulative findings from the addiction field, leads us to conclude that compounds that target 5-HT_2A_R signaling are excellent candidates for the modification of drug seeking behavior. Because psilocybin is a 5-HT_2A_R agonist, these results suggest that modulation of 5-HT_2A_R signaling may be critical for maintaining abstinence from opioid seeking or for reducing opioid-related craving behavior.

Opioids trigger a proinflammatory response ^66, 67^ and published studies have demonstrated the utility of reducing inflammatory signaling to inhibit drug seeking behaviors ^68, 69^. Inhibition of toll-like receptor 4 (TLR4), which is associated with opioid addiction and can activate TNF-α/NF-kB pathways, inhibits opioid seeking behaviors ^70–72^. Intriguingly, activation of 5-HT_2A_R with the agonist (*R*)-DOI abolished the inflammatory effects mediated by TNF-α systemic administration in mice ^27^. Moreover, opioid-induced TNF-α dysregulation was reversed in peripheral blood samples from OUD patients treated with the medications buprenorphine or methadone ^68^. Hence, targeting this receptor may be a viable option for modulation of inflammatory signaling pathways, such as those that are impacted by chronic opioid exposure, as indicated by our observation of a negative relationship between heroin relapse behavior and expression of 8 inflammatory cytokines or chemokines in the brain when animals were pretreated with psilocybin.

Surprisingly, we did not observe significant changes in gene expression across the qPCR panel, which included numerous cytokines, chemokines, and their respective receptors, that mirrored the RNA sequencing data when psilocybin was administered 24 hr prior to a relapse test. This lack of significant variation may be attributed to the dynamic nature and rapid turnover of chemokines and cytokines, coupled with the short half-life of these inflammatory factors ^73, 74^. Moreover, it is important to consider that our analysis was confined to mRNA levels rather than protein levels. Given the dynamic and pulsatile behavior of these inflammatory factors, it is plausible that their mRNA levels had returned to baseline after 24 hours, which might not directly reflect changes at the protein level. Future studies could benefit from examining the protein pool to provide deeper insights into these discrepancies. The consistency in behavioral outcomes between the 4-hour and 24-hour pretreatment groups might be explained by the possibility that protein levels, which can persist longer than their mRNA transcripts, differ from the mRNA profiles. Therefore, while mRNA levels provide valuable information, they may not fully capture the changes at the protein level that are crucial for mediating the behavioral effects observed. Alternatively, the transcriptomic changes observed at the inflammatory factor level could affect pathways not directly assessed in our current study, but which play critical roles in modulating behavior. These pathways may include, but are not limited to, those involved in synaptic plasticity, and neurotransmitter signaling. While our study focused on specific markers of inflammation, the broader impact of these transcriptomic changes may extend to these critical pathways, influencing behavioral outcomes in ways not captured by our initial analysis.

However, we did observe regulation in the IL-17a pathway in both PFC qPCR and sequencing datasets of psilocybin-treated rats: In animals that displayed psilocybin-mediated inhibition of heroin seeking following a 4 hr acute psilocybin pretreatment, 3 mg/kg psilocybin decreased PFC *Il17a;* while *Il17ra*, the receptor for IL-17a, was downregulated following a single exposure to psilocybin 24 hr prior in our RNA sequencing study. Alcohol-dependent mice treated with anti-IL-17a antibodies reduced their voluntary alcohol intake ^75^, suggesting that modulation of IL-17a may impact drug seeking behavior. Additionally, IL-17a is proinflammatory and was reported upregulated in the blood of mice following long-access exposure to fentanyl vapor ^76^, as well as in the PFC after methamphetamine exposure ^77^. Treatment with a purinergic P2X7 receptor antagonist, which blocks pro-inflammatory signaling ^78^, inhibits methamphetamine-induced upregulation of IL-17a in the PFC, as well as methamphetamine-induced conditioned place preference ^77^. We conducted an additional experiment showing that selective inhibition of IL-17a signaling in the PFC before a relapse test significantly reduced heroin-seeking behavior in animals after 21 days of forced abstinence from heroin SA. This finding corroborates previous research suggesting that targeting IL-17a signaling could be an effective strategy for reducing drug-seeking behaviors. Additionally, this experiment supports our hypothesis that modulating inflammatory pathways in the PFC may be one of the mechanisms through which psilocybin achieves its therapeutic effects, particularly in diminishing drug-seeking behaviors. Therefore, studying other inflammatory pathways could also be helpful. For example, psilocybin also increased NAc *Tnfrsf11b,* which encodes the Opg protein. This protein is elevated by anti-inflammatory drugs ^79–81^, and has been studied for neuroprotective and inflammatory effects ^82^, although not for drug seeking behavior. Future studies exploring downstream signaling consequences of psilocybin will provide additional pathways that may be therapeutic targets for reduction of opioid seeking.

In this study, we have directed our molecular investigations downstream of psilocybin exposure towards inflammatory cytokines and chemokines, driven by emerging evidence that emphasizes reports of opioid-induced regulation of inflammatory modulators as a major neuroadaptation observed in the molecular foundations of OUD ^30, 83^. Additionally, multiple lines of evidence demonstrate that inflammation significantly impacts the regulation of synaptic plasticity^84–86^ and it is posited that psychedelics such as psilocybin may induce long-lasting therapeutic effects by altering the nature of neural circuits through neuronal plasticity. Psychedelic compounds may facilitate structural changes within the nervous system, rendering it more receptive to environmental stimuli. This heightened receptivity may enable extensive modifications at the synaptic, circuit, and behavioral levels, through induction of a specific form of synaptic plasticity termed metaplasticity, which supports the restoration and stabilization of healthy neural function^87, 88^. Psilocybin can modulate neuroimmune factors and this interaction could represent one of the mechanisms through which psychedelics facilitate changes in synaptic plasticity, potentially leading to durable therapeutic effects. Intriguingly, this positions psychedelics as potential therapeutic agents in OUD by potentially resetting maladaptive neural circuits and reducing neuroinflammation. This dual action highlights the innovative potential of psychedelics in redefining treatments for addiction and other psychiatric disorders where neuroinflammation and synaptic dysfunction play critical roles. As such, further research into the interplay between psychedelic-induced neuroplasticity and neuroimmune modulation is crucial for developing targeted therapies that harness these complex biological interactions.

While these exciting findings demonstrate psilocybin reduces heroin relapse, limitations of the study include use of only male animals, given that sex-specific effects of psilocybin on mouse ethanol drinking ^46^ and responses to 5-HT_2A_R psychedelics have been reported ^89–91^. In our initial investigation, we employed a subchronic treatment with high doses of psilocybin, an approach that may lack translational validity, to enable direct comparisons with both the microdosing protocol and the ketanserin antagonist treatment. Nevertheless, we subsequently replicated the experiments using a single high-dose psilocybin administration at two distinct time points. The consistent results from these tests, even at the later time point (24 hr), suggest that psilocybin’s positive effects on drug-seeking behavior may be long lasting. Secondly, for the initial molecular studies, we used naive animals, avoiding any variables associated with voluntary heroin use or withdrawal. This approach provided a clear baseline for pharmacological comparisons, yet it also attenuated the translational potential of our transcriptomic observations in the context of relapse-tested animals with a history of heroin self-administration. To address this limitation, we expanded our analysis focusing on key pathways revealed by RNA-seq in samples from animals that underwent relapse tests at different time points. Although we have initially focused our study on the transcriptional alterations in only the PFC and NAc following psilocybin exposure, psilocybin has been shown to impact many other brain regions ^92^. We acknowledge that the sample size used in Figure 1 to investigate the impact of 0.1 mg/kg and 3 mg/kg psilocybin on heroin SA may be underpowered to detect small differences and future studies replicating our data in larger cohorts are undoubtedly necessary. While considering these potential limitations, the findings indicated that inflammatory mediators may play a role in the psilocybin-induced reduction of heroin-seeking behaviors. Future studies into putative sex-specific responses to psilocybin during heroin relapse are warranted. Finally, psilocybin can also activate other receptor systems, such as 5-HT_2C_R, 5-HT_1A_R and TrkB ^61, 63, 93^, and whether the effects of psilocybin on heroin relapse are mediated entirely through 5-HT_2A_R, or if non-5-HT_2A_R mechanisms also contribute to psilocybin-mediated reduction of heroin relapse is currently unknown. Ketanserin may also function as a 5-HT_2C_R antagonist and future studies with additional 5-HT_2A_R/5-HT_2C_R antagonists in combination with psilocybin are needed. Irrespective of these potential limitations, our data suggests that psilocybin has a promising role in modulating heroin-seeking behavior, potentially offering a novel therapeutic avenue for the treatment of opioid addiction.

## Acknowledgements

Heroin was provided by the National Institute on Drug Abuse drug supply program. We thank all members of the Daws lab for helpful discussion on the interpretation of data.

## Author contributions

Conceptualization, S.D.; methodology, S.D., G.F.; formal analysis, S.D., G.F., K.D.; data collection, G.F., M.Z.; writing- original draft preparation, S.D., G.F.; writing- review and editing: S.D., G.F., K.D., M.Z.; supervision, S.D.; project administration, S.D.; funding acquisition, S.D. All authors have read and agreed to the published version of the manuscript.

