## Supplemental Information for "Psilocybin reduces heroin seeking behavior and modulates inflammatory gene expression in the nucleus accumbens and prefrontal cortex of male rats"

**Supplemental methods:**

*Jugular vein catheterization and heroin self-administration (SA)*

Jugular vein catheterization and heroin SA were performed as previously described (1, 2). After surgery, rats received antibiotics (Cefazolin 10 mg/kg, intravenously) for five consecutive days and recovered for one week before beginning behavioral testing. Catheters were flushed with heparin solution (10 USP/ml) daily to maintain patency. Rats underwent a patency test with propofol (1%) prior to drug self-administration (SA). Heroin SA and relapse tests were performed in operant chambers (29.5 × 32.5 × 23.5 cm, Med Associates, Fairfax, VT, USA). Briefly, rats self-administered 0.075 mg/kg/infusion heroin under a fixed ratio (FR) 1 schedule for 10-11 days in 6-hour daily sessions. Infusion of heroin solution was paired with the presentation of a 65 db, 2.9 kHz acoustic cue and a stimulus light placed above the active lever; the inactive lever did not produce any response. Rats were excluded from the study if they did not make at least 10 infusions per day for the last 4 days of SA or did not show a 2-fold preference for the active versus inactive lever. The syringe volume was noted before and after each heroin session and compared to the number of infusions to ensure the correct volume of heroin was administered. For data in Figures 1-4, animals were not 're-randomized' after the SA testing, prior to the relapse tests. Animals received the same psilocybin, ketanserin or volinanserin dose prior to the relapse test that they had received prior, during days 9-11 of training.

*Locomotor assessment*

Assessment of locomotor activity was performed as previously described (3). Briefly, rats were individually placed into transparent plastic chambers (45 cm × 20 cm × 20 cm) that were set within metal frames containing 16 infrared light emitters and detectors. The intervening space between the beams was 2.5 cm and beam height were 4.5 cm. A computer interface with a dedicated software (Digiscan DMicro system, Accuscan, Inc., Columbus, OH) recorded the number of photocell beam breaks.

*Gene expression array*

Total RNA was resuspended in RNase free water and concentration was evaluated with a Qubit 3.0 Fluorometer using the Qubit RNA High Sensitivity Assay buffer (Invitrogen, Carlsbad, California, USA). For measurement of gene expression with the RT² Profiler™ PCR Array, 0.5 µg of total RNA was reverse transcribed with RT² First Strand Kit (Qiagen, Hilden, Germany) in a MiniAmp Thermal Cycler (Thermo Fisher Scientific) according to manufacturer's instructions. cDNA amplification to measure inflammatory cytokines and receptors was performed using a Quantstudio3 qPCR machine (Thermo Fisher Scientific, Waltham, MA, USA) according to manufacturer's instructions. Replicates were excluded from qPCR analysis if a Ct value ≥ 35. Differential expression analyses were performed using the $\Delta\Delta$ct method of analysis (4) and statistics were performed on $\Delta\Delta$ct values prior to log transformation into fold change. All animals that completed the relapse test in Fig. 6A-I were euthanized within 1 hour of the conclusion of the behavioral experiments, and RNA extracted for qPCR array analysis.

*RNA sequencing and bioinformatic analysis*

RNA sequencing of PFC tissue following a single exposure to psilocybin or ketanserin was performed on 400 ng of RNA per sample, with 4 biological replicate samples per treatment group. Paired-end libraries were generated from total RNA using poly-T oligo-attached magnetic beads. First strand cDNA was created with random hexamer primers, followed by second strand cDNA synthesis using dUTP. End repair, A-tailing, adapter ligation, size selection, amplification and purification of libraries were performed using proprietary reagents at Novogene. The library was analyzed for quality control with Qubit, PCR, and bioanalysis for size distribution detection. Quantified libraries were pooled and sequenced on the Illumina NovaSeq 6000 platform. Raw reads were processed to removed reads containing adapters, poly-N or low quality. Hisat2 v2.0.5 (5) was used to map reads to the rn6 reference genome and StringTie (v1.3.3b) was used to assemble mapped reads (6). featureCounts v1.5.0-p3 was used to count reads mapped for each gene (7) and calculate the FPKM, expected number of Fragments Per Kilobase of transcript sequence per Millions base pairs sequenced. DESeq2 R package (1.20.0) (8) was used to determine differential expression between two groups, which uses a model based on the negative binomial distribution. Genes with an average expression value of less than 5 for both comparison groups were excluded from the analysis. Genes with >30% fold change and p-value ≤ 0.05 were considered statistically significant. The Database for Annotation, Visualization, and Integrated Discovery (DAVID version 6.8; https://david.ncifcrf.gov) was used for gene ontology and enrichment analysis. Rank-Rank Hypergeometric Overlap (RRHO) (9) plots were generated to compare the overlap between transcriptomic signatures for two distinct experimental conditions (e.g., Vehicle-Psilocybin 3mg/kg and Vehicle-Ketanserin 3mg/kg compares DEG list from Psilocybin 3mg/kg vs Vehicle against the DEG list from Ketanserin 3mg/kg vs Vehicle). The stratified RRHO plots were generated using the RRHO2 package that was optimized by Li Shen lab (github.com/shenlab-sinai/RRHO2). To generate alluvial plots, genes in each experimental condition were categorized as upregulated (up; p-value ≤ 0.05 and Log2FoldChange ≥ 0.378), downregulated (down; p-value value ≤ 0.05 and Log2FoldChange ≤ -0.378) or not significant (ns). Common regulatory patterns across conditions were mapped using the ggalluvial package in R (version 0.12.5). GeneOverlap was used to determine statistical significance of two transcriptome datasets using the Fisher's exact test. To perform the Fisher's exact test, gene lists of significantly differentially regulated genes for each condition were prepared (p-value ≤ 0.05 and Log2FoldChange ≥ 0.378 or Log2FoldChange ≤ -0.378). The gene lists for each condition were compared using the GeneOverlap package in R Bioconductor, version 1.36.0.

*Primer List*

The following gene expression assays were purchased from Integrated DNA Technologies and used in the study:

*Fos*: forward primer TTG GCA CTA GAG ACG GAC A, reverse primer CAG CCT TTC CTA CTA CCA TTC C, and probe /56-FAM/CTG TCA ACA /ZEN/CAC AGG ACT TTT GCG C/3IABkFQ/

*ActB*: forward primer GGC ATA GAG GTC TTT ACG GAT G, reverse primer TCA CTA TCG GCA ATG AGC G, and probe /56-FAM/TCC TGG GTA /ZEN/TGG AAT CCT GTG GC/3IABkFQ/

*Gapdh*: forward primer GTA ACC AGG CGT CCG ATA C, reverse primer TCT CTG CTC CTC CCT GTT C, and probe /56-FAM/CAC ACC GAC /ZEN/CTT CAC CAT CTT GTC T/3IABkFQ/

**Supplemental figure 1: Psilocybin does not increase heroin self-administration.**

(A-D) Individual animal responses during SA to each dosage of psilocybin or vehicle. Each line represents one animal. Displayed are active lever responses averaged prior to (Pre; days 6-8) or after (Post; days 9-11) psilocybin pretreatment.

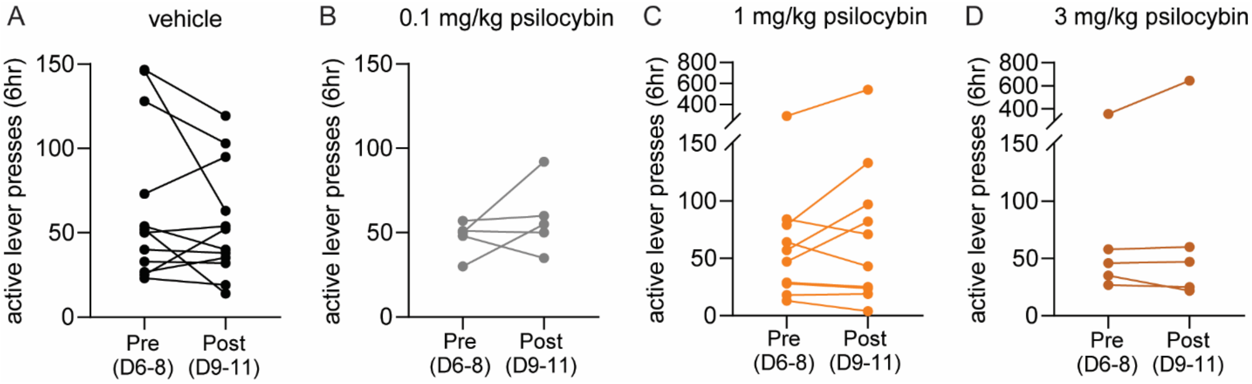

**Supplemental figure 2: Ketanserin increases heroin self-administration and intake at the highest dose examined.**

(A-D) Individual animal responses during SA to each dosage of ketanserin or vehicle. Each line represents one animal. Displayed are active lever responses averaged prior to (Pre; days 6-8) or after (Post; days 9-11) ketanserin pretreatment. * paired t-test, $p<0.05$. (E) Infusions made by 8 individual animals during heroin SA pre (day 8, black line) or post (day 11, red line) treatment with 3.0 mg/kg ketanserin. Each downward line represents a drug infusion.

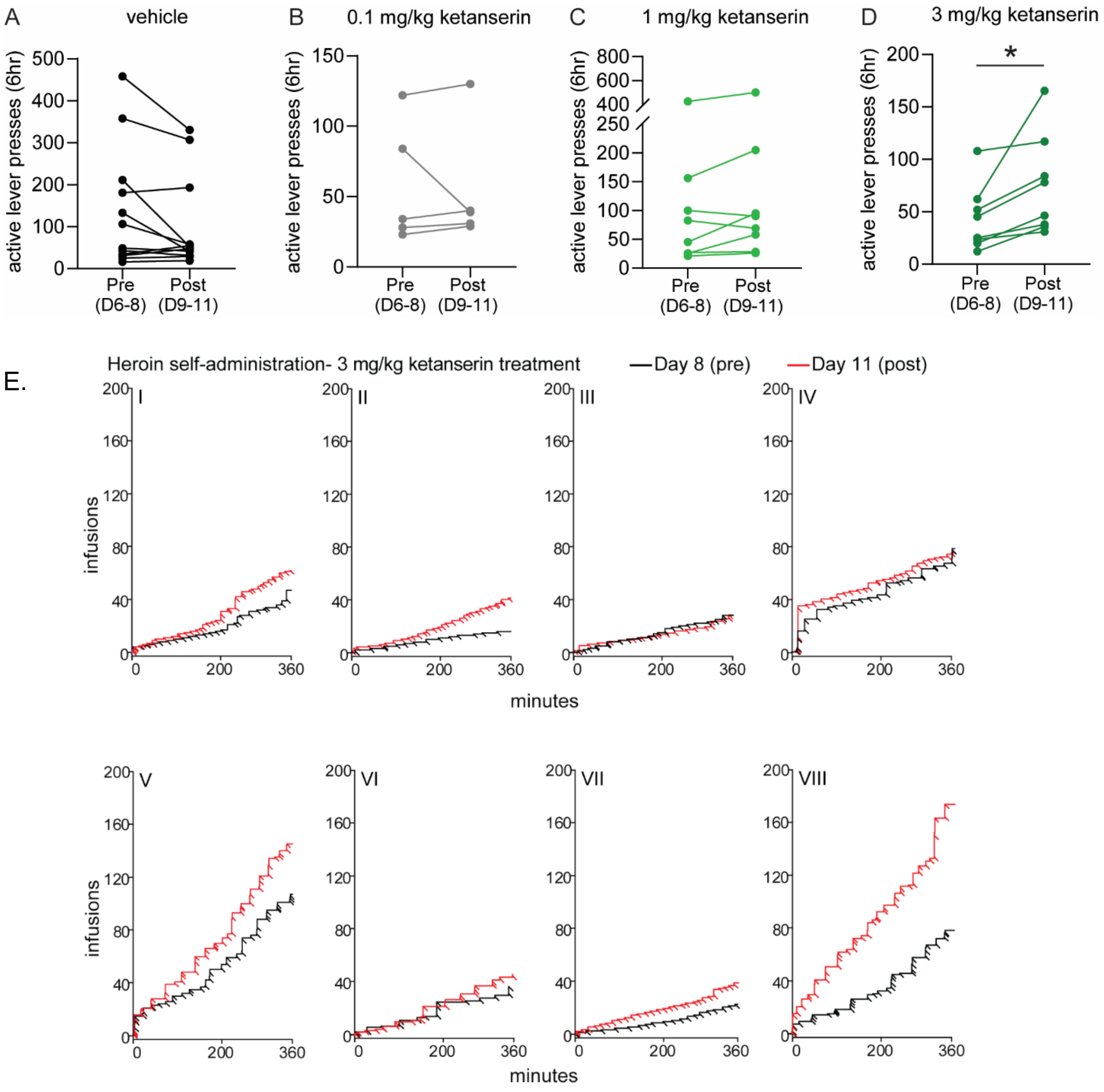

**Supplemental Figure 3: Treatment with ketanserin or psilocybin does not impact general locomotor activity.**

Following treatments with psilocybin or ketanserin described in Fig 1-2, a subset of animals was tested for locomotor activity 2 weeks after the completion of relapse test 3 (R3). (A) Animals received a single IP injection of vehicle or 3 mg/kg ketanserin. 15 minutes later, locomotor activity was assessed for 30 min in an activity chamber. (B-C) Animals received a single injection of vehicle or 3 mg/kg psilocybin (B), or 1 mg/kg psilocybin (C). 4 hr later, locomotor activity was assessed for 30 min in an activity chamber. Error bars indicate standard error of the mean. *n*=5-6/group.

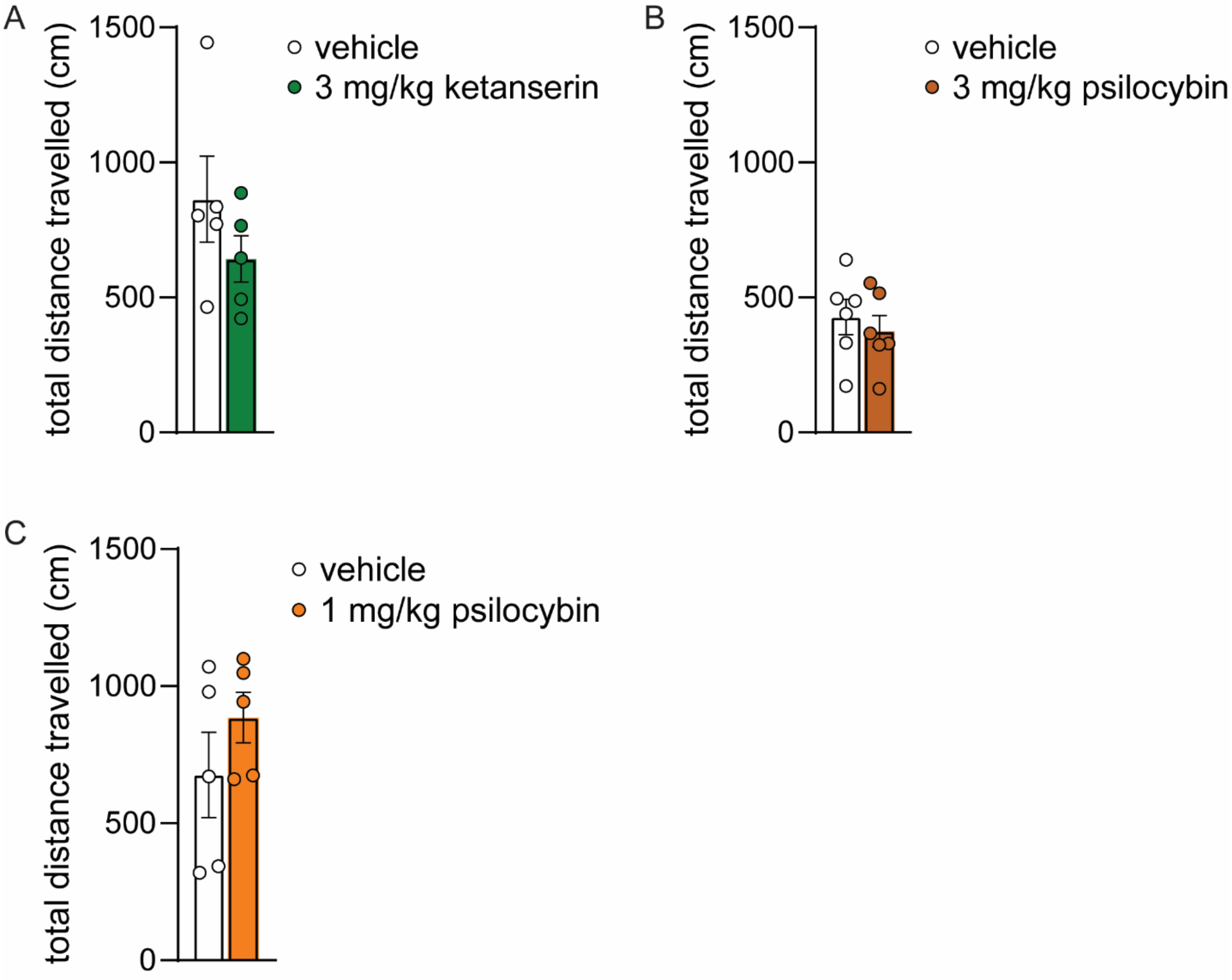

**Supplemental Figure 4: Alluvial plots of gene expression patterns in the PFC following treatment with psilocybin.** Alluvial plots representing clusters of genes that exhibit specific patterns of gene expression through: (A) 3mg/kg psilocybin, 3mg/kg psilocybin + 3mg/kg ketanserin, and 3mg/kg ketanserin; (B) 1mg/kg psilocybin, 3mg/kg psilocybin, 3mg/kg psilocybin + 3mg/kg ketanserin; (C) 1mg/kg psilocybin, 3mg/kg psilocybin, 3mg/kg ketanserin. On the x-axis, genes are stratified for every experimental condition, to show the number of genes on the y-axis corresponding to each cluster, as represented by the color-coding. Up- upregulated, down- downregulated, ns – not significant.

A.

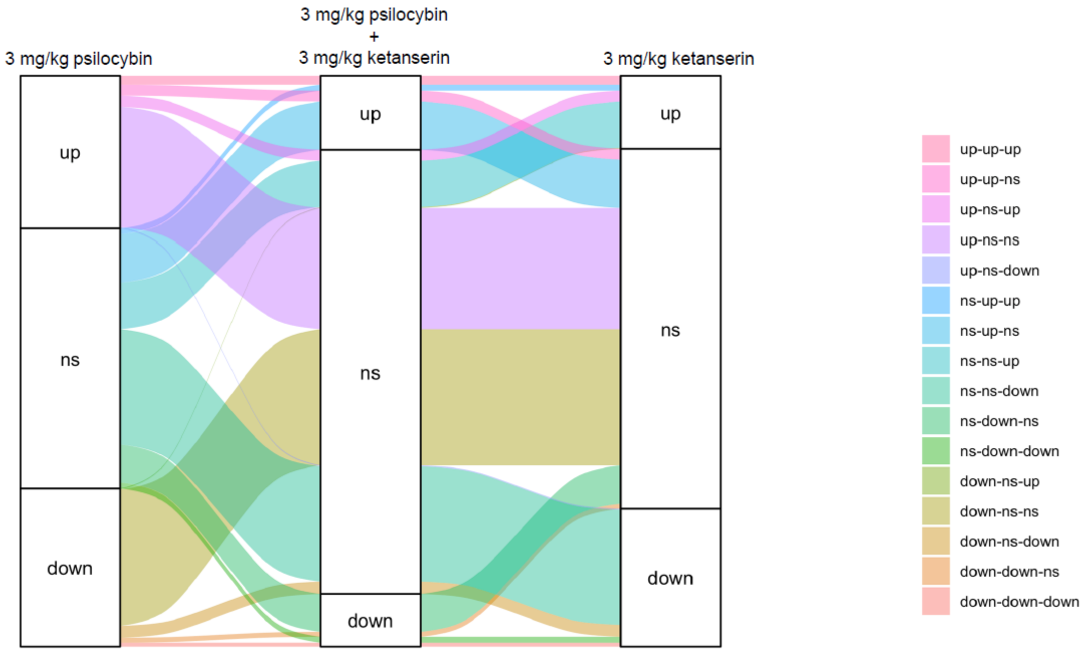

B.

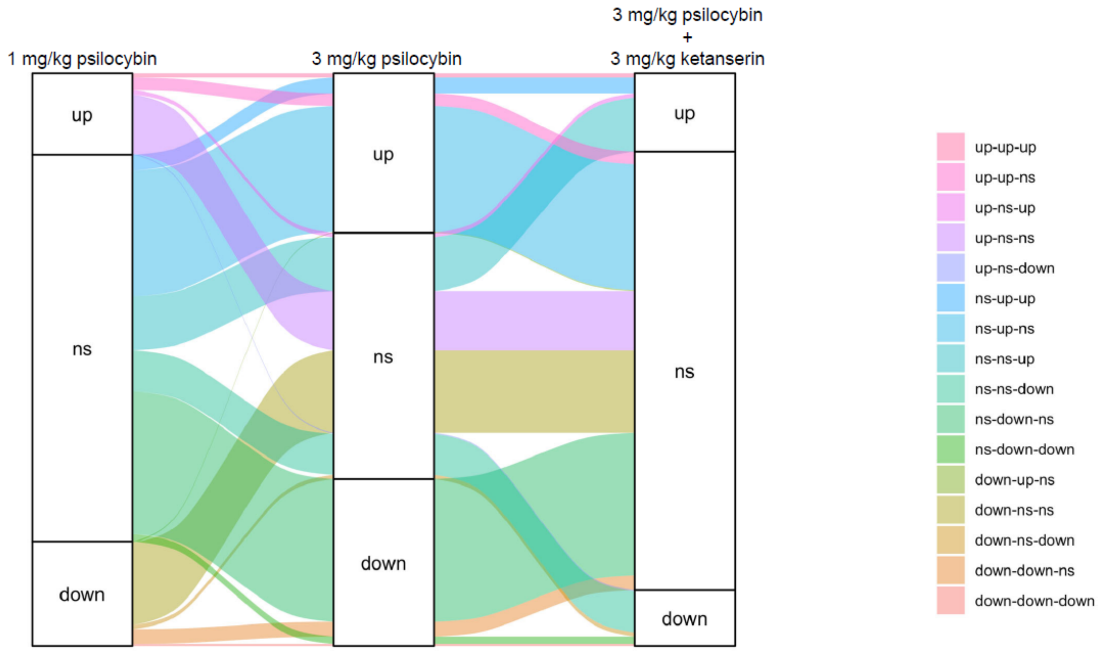
1 mg/kg psilocybin
3 mg/kg psilocybin
3 mg/kg psilocybin
+
3 mg/kg ketanserin
up
ns
down
up
ns
down
up
ns
down
up-up-up
up-up-ns
up-ns-up
up-ns-ns
up-ns-down
ns-up-up
ns-up-ns
ns-ns-up
ns-ns-down
ns-down-ns
ns-down-down
down-up-ns
down-ns-ns
down-ns-down
down-down-ns
down-down-down

C.

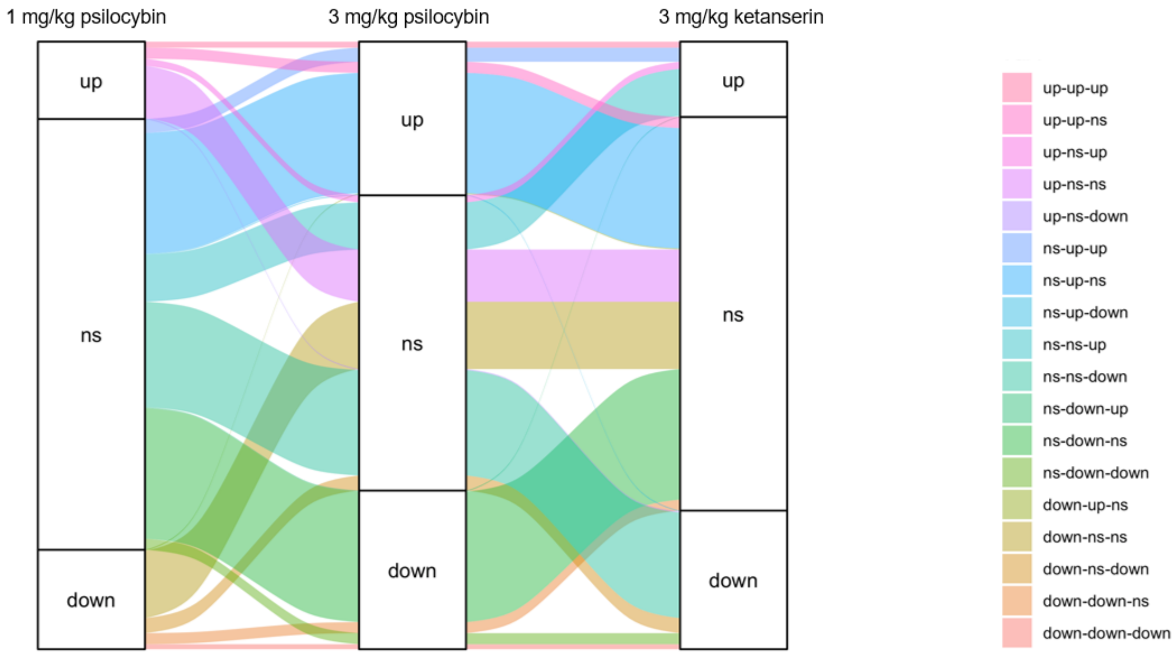
1 mg/kg psilocybin
3 mg/kg psilocybin
3 mg/kg ketanserin
up
ns
down
up
ns
down
up
ns
down
up-up-up
up-up-ns
up-ns-up
up-ns-ns
up-ns-down
ns-up-up
ns-up-ns
ns-up-down
ns-ns-up
ns-ns-down
ns-down-up
ns-down-ns
ns-down-down
down-up-ns
down-ns-ns
down-ns-down
down-down-ns
down-down-down

**Supplemental Figure 5: Animals that receive a single vehicle injection prior to a relapse test display similar relapse behavior.**

Rats underwent heroin SA followed by forced abstinence, as described in Figure 5. Rats received a single vehicle injection of saline either 4hr or 24hr prior to a relapse test. Shown are active and inactive lever responses for both groups of vehicle-treated rats. SA and relapse data for vehicle-treated rats are collapsed into a single group in Figure 5. Error bars indicate standard error of the mean. *n*=7-8/group.

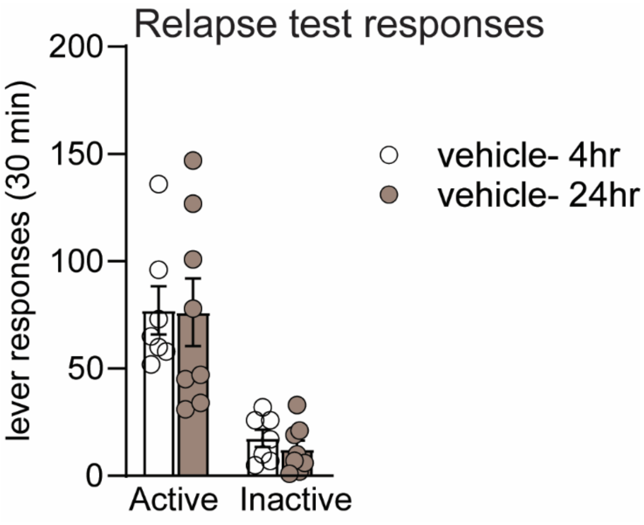

**Supplemental Figure 6: Psilocybin regulates gene expression of inflammatory cytokine and chemokines in the NAc during inhibition of heroin seeking.**

(A) Volcano plots depicting inflammatory cytokine and chemokine genes differentially expressed in the NAc of rats that were treated with 3.0 mg/kg psilocybin 4hr prior to a relapse test, vs vehicle treated animals. Horizontal dotted line denotes p value of 0.05. Vertical dotted line denotes fold change of +/- 30%. Grey circles represent genes that were not significantly altered. Red dots represent genes that meet criteria for significance. (B) Results of correlation analysis of relapse behavior of rats (from Figure 5A) with inflammatory cytokine or chemokine gene expression data for the NAc. Italicized, bolded values were statistically significant.

A

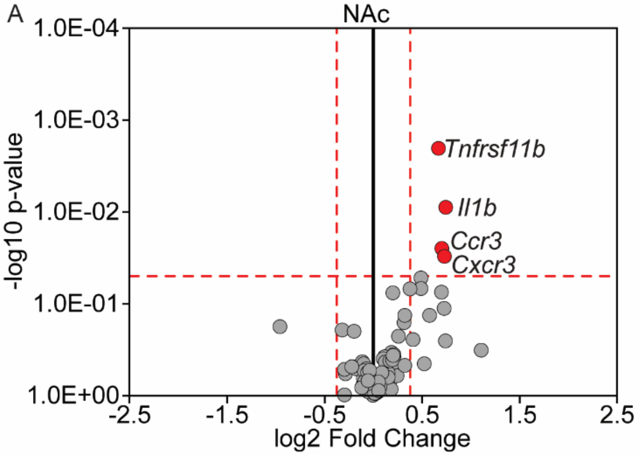


B

| Nac | heroin SA + saline | | heroin SA + 3 mg/kg psilocybin | |
|---|---|---|---|---|
| | Pearson r | *p*-value | Pearson r | *p*-value |
| *Ccl7* | -0.148 | 0.752 | ***-0.891*** | ***0.007*** |
| *Ccr10* | -0.539 | 0.211 | ***-0.774*** | ***0.041*** |
| *Ccr2* | -0.334 | 0.465 | ***-0.837*** | ***0.019*** |
| *Faslg* | -0.396 | 0.379 | ***-0.768*** | ***0.044*** |
| *Il33* | -0.700 | 0.080 | ***-0.786*** | ***0.036*** |
| *Pf4* | -0.132 | 0.778 | ***-0.885*** | ***0.008*** |
| *Tnfrsf11b* | 0.068 | 0.885 | ***-0.803*** | ***0.030*** |
